# A Network-based Transcriptomic Landscape of HepG2 cells to Uncover Causal Gene Cytotoxicity Interactions Underlying Drug-Induced Liver Injury

**DOI:** 10.1101/2023.01.16.524182

**Authors:** Lukas S. Wijaya, Attila Gabor, Iris E. Pot, Luca van de Have, Julio Saez-Rodriguez, James L. Stevens, Sylvia E. Le Dévédec, Giulia Callegaro, Bob van de Water

**Affiliations:** Leiden Academic Centre for Drug Research (LACDR), Leiden University, Leiden, The Netherlands; Heidelberg University, Faculty of Medicine, and Heidelberg University Hospital. Institute for Computational Biomedicine, 69120 Heidelberg, Germany

## Abstract

Drug-induced liver injury (DILI) remains the main reason of drug development attritions largely due to poor mechanistic understanding. Toxicogenomics to interrogate the mechanism of DILI has been broadly performed. Gene network-based transcriptome analysis is a bioinformatics approach that potentially contributes to improving mechanistic interpretation of toxicogenomics data. In this current study, we performed an extensive concentration time course response-toxicogenomics study in the HepG2 cell line exposed to various DILI compounds, reference compounds for stress response pathways, cytokine receptors, and growth factor receptors. We established > 500 conditions subjected to whole transcriptome targeted RNA sequences and applied weighted gene co-regulated network analysis (WGCNA) to the transcriptomics data followed by identification of gene networks (modules) that were strongly modulated upon the exposure of DILI compounds. Preservation analysis on the module responses of HepG2 and PHH demonstrated highly preserved adaptive stress responses gene networks. We correlated gene network with cell death as the progressive cellular outcomes. Causality of the target genes of these modules was evaluated using RNA interference validation experiments. We identified that *GTPBP2, HSPA1B, IRF1, SIRT1* and *TSC22D3* exhibited strong causality towards cell death. Altogether, we demonstrate the application of large transcriptome datasets combined with network-based analysis and biological validation to uncover the candidate determinants of DILI.

## Introduction

Drug-induced liver injury (DILI) remains a worldwide health problem, representing 21% of adverse drug reactions^1,2^. Additionally, DILI is also the main cause for drug attritions in both clinical and preclinical phases^3^. Multiple studies have been conducted with the goal to improve the understanding of the mechanisms of DILI. For example, one recent review reported multiple cellular responses that could lead to clinical manifestation of DILI: mitochondrial impairment, inhibition of biliary efflux, lysosomal impairment, reactive metabolites, endoplasmic reticulum stress, and immune system activation^4^. Despite recent advances a more holistic understanding of the cellular events underpinning DILI outcomes and their causal relationship with adverse events starting from cell to more complex tissue and organ responses in an AOP framework is yet to be developed. In this case, the lack of the reliable data interpretation describing the causality between the cellular responses and cellular outcomes in DILI episodes becomes a limitation hurdling the exploration.

Transcriptomic approaches are promising tools to achieve a more holistic understanding of the mechanisms of DILI as well as the prediction of its occurrence^5–7^. For example, an extensive *in vitro* study has been performed in HepaRG cell lines exposed to >1,000 chemicals from the ToxCast library assessing the expression of almost 100 different genes. Despite this small gene set, this study identified transcriptomics signatures related to molecular initiating events of these compounds^8^. In other contexts, transcriptomic profiling has been reported to have higher predictability towards cellular outcomes compared to a sole interpretation from the activation of transcription factors^9^. However, interpreting transcriptomic data can be challenging due to high dimensionality of the data resulting to low signal-to-noise and high variation^10^. Enrichment approaches such as pathway enrichment analysis and network-based analysis help reduce the dimensionality of high content transcriptomic datasets problem^11^. The network-based analysis is becoming a promising approach to study complex biological responses. Barabási et al. have introduced the notion that network-based approach improve the understanding of disease pathways and the identification of potential drug target and biomarkers^12^. Such approaches could be applied to DILI-related toxicogenomics data to advance the mechanistic understanding of DILI.

Weighted gene co-regulated network analysis (WGCNA) is one method to perform network-based analysis^13^. This method clusters genes based on their co-expression into specific networks called modules. Each module is scored (eigengene scores) based on the expression values of the composing genes, thus indicating an overall value of the module activity (induction or repression). In order to increase the biological meaning of the modules, each module can be enriched to certain annotations linked to the cellular responses. To date, a few studies have applied WGCNA on toxicogenomics datasets. Sutherland et al, derived WGCNA modules from the *in vivo* transcriptomic data in rats based on the Drug Matrix dataset and demonstrated that modules facilitated to unravel mechanisms of DILI in rats and to define molecular processes that underlie pre-clinical outcomes, such as histopathology and clinical chemistry^14^. Callegaro, et al developed a WGCNA approach usingcultured primary human hepatocytes (PHH) from the TG-GATEs dataset. WGCNA modules were able to capture the hepatocellular events related to DILI and furthermore identified the cellular events that were conserved across species^15^. So far, an attempt to systematically define the causality between gene network perturbation and onset of toxicity is lacking. While PHH represent the gold standard for *in vitro* DILI testing, PHH have inherent limitations for genetic manipulation to uncover mechanisms. In contrast, the hepatocarcinoma HepG2 cell line, which is widely used for early DILI screening and showed relatively close resemblance to cytotoxicity responses of PHH^16^, allows mechanistic functional genomics studies. However, so far large toxicogenomics datasets of HepG2 that allow WGCNA application are missing. Moreover, the knowledge on the preserved cellular responses linked to progressive cellular outcomes between PHH and HepG2 is undefined. The utilization of robust, cheap and fast liver test systems for WGCNA application will facilitate the experimental validation of causal relationships between module activation and cellular outcomes.

In this study, we established a comprehensive toxicogenomics dataset with the HepG2 cell line based on exposure to over 500 different conditions. This involved DILI compounds, reference compounds that activate specific stress response pathways, cytokines and growth factors, and reference compounds^17^. Cells were treated with the substances with up to six different concentrations and high throughput TempO-Seq targeted RNA sequencing was performed at 4, 8, and 24 hours. We utilized the WGCNA approach to capture the cellular events in the context of DILI compound treatment and mapped the highly perturbed cellular responses upon exposure of the DILI compounds. We performed preservation analysis with PHH and defined modules that are associated with DILI compound-induced cell death. Finally, we validated the causality between the high cell death-correlated modules with the cell death outcomes. From the 67 highly upregulated genes inside correlated modules we found *GTPBP2, HSPA1B, IRF1, SIRT1* and *TSC22D3* to impact on cell death onset (Figure 1A). Altogether, we established a novel quantitative network-based approach for the assessment of HepG2 toxicogenomics data that enables improved mechanistic understanding of mode-of-action of DILI compounds and novel drug candidates.

**Figure 1.**
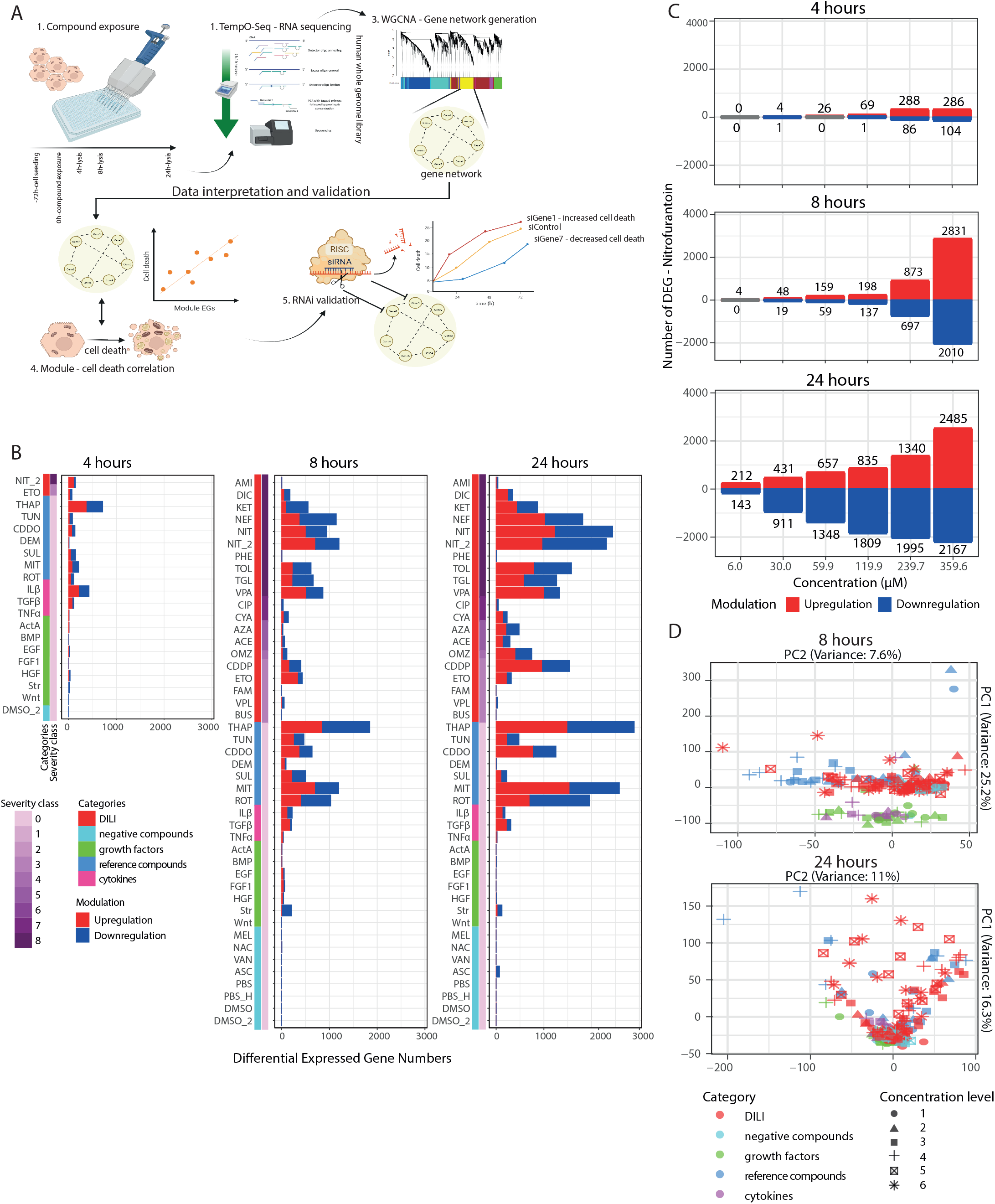
Temporal TempO-Seq whole transcriptome targeted RNA sequencing of HepG2 cells. (A) The experimental overview of the study showing the schematic time line of the cell culture processes to the high throughput transcriptomic (HTTr) method application. The cells were seeded in 384 well plate 72 hours before the exposure to the compounds. The cells were lysed and then the lysates were collected for HTTr processes with BioSpyder technology at 4 hours, 8 hours, and 24 hours. The lysates were then subjected to the TempO-Seq high throughput RNA sequencing technology deploying the whole genome library. The log2 fold change values were used to generate gene networks with WGCNA approach. Correlation with the external trait-cell death were performed from the previously generated cell death data^17^. The causal relation of gene memberships from the high cell death correlated-modules with the cell death occurrence were determined using RNAi method. (B) The differential expressed gene (DEG) numbers of each tested compound in every time point. Each plot shows the aggregated sum values from all the concentration. The color bars on the rows show compound categories and severity class. In the bars, the blue color indicates the downregulated DEGs and the red color shows the upregulated DEGs. The threshold of the DEGs is set with adjusted p-value < 0.01 and log2 fold change > [0.1]. (C) The differential expression gene numbers of the cells exposed to nitrofurantoin from the lowest dose level (1) until the highest dose level (6) in every time point. The red color of the plot indicates upregulated DEGs and blue color indicates downregulated DEGs. The threshold of the DEGs is set with adjusted p-value < 0.01 and log2 fold change > [0.1]. (D) The PCA plots derived from the log2 fold change values, per time point. Every dot of the plots indicates the position of each sample in the plot where the colors indicate the compound category and the shapes represent the dose levels.

## Material and Method

### 2.1 Chemical and Reagents

All chemicals were purchased from Sigma-Aldrich – The Netherlands; except for cisplatin (Ebewe – The Netherlands) and nefazodone (Sequoia Research Products – Pangbourne, United Kingdom). All compounds were dissolved in DMSO; except for mitomycin-C (DMSO-PBS) and N-acetylcysteine (PBS) and for cisplatin which was already manufactured as a solution. All compounds in DMSO were maintained as 500-fold stock such that the final exposure did not exceed 0.2% v/v DMSO. The cytokines and growth factors are dissolved according to the manufacture protocols. TNFα was purchased from R&D System (Abingdon, United Kingdom), TGFβ was purchased from Immunotools (Friesoythe, Germany), ActivinA, BMP4, hHGF, hFGFi, hIL-1β, and WNT3a were purchased from PeproTech (London, United Kingdom), human EGF was purchased from Sigma-Aldrich (The Netherlands). PowerUp SYBR green real time PCR master mix was purchased from ThermoFisher.

### 2.2 Cell line

Human hepatoma (HepG2) cells were purchased from ATCC - Germany (clone HB8065) and maintained in DMEM high glucose (Fisher Scientific – Bleiswijk, The Netherland) supplemented with 10% (v/v) FBS (Fisher Scientific-Bleiswijk, The Netherlands), 250 U/ml penicillin and 25 μg/ml streptomycin (Fisher Scientific – Bleiswijk, The Netherlands) in humidified atmosphere at 37 degrees Celsius and 5% CO2 /air mixture. The cells were used between passage 14 and 20. The cells were seeded in Greiner black μ-clear 384 well plates, at 8000 cells per well for the exposure experiment. Unless differently mentioned, we use wild type HepG2 for the experiments.

### 2.3 Compound exposure and cell lysis

The exposure of the compounds and the RNA sequencing was performed at different moments and referred to as batch 1 and batch 2. The 1^st^ batch was performed for the all DILI compounds and the 2^nd^ batch was performed for all cellular stress response reference compounds, cytokines, and growth factors; nitrofurantoin was included in both batches at the same concentrations to assess inter-batch variation. The DILI severity of the compounds was adapted from the FDA^18^. At the day of exposure (three days after seeding), the compounds were diluted in the culturing medium to meet the final concentration was as indicated in Table 1. The plates were incubated in humidified atmosphere at 37 degrees Celsius and 5% CO_2_ /air mixture. At the designated time points (4 hours –1^st^ and 2^nd^ batch, 8 hours, and 24 hours – 2^nd^ batch) the cells were lysed. The exposure medium was aspirated and the cells were washed with PBS 1 time followed by addition of 25 μL 1x BNN lysis buffer (BioClavis, Glasgow, Scotland) diluted in PBS. The plates were incubated for 15 minutes at room temperature to enhance the lysis processes. After 15 minutes incubation, the plates were sealed with the aluminium seal and stored at -80 degree Celsius before sequencing. The plates were shipped to BioClavis where the RNA sequencing was performed. All exposures were performed three times independently to cover biological variability.

**Table 1.** List of the tested compounds with the supporting information. The severity of the DILI compounds are defined from a previous study^18^.

| Compound list (concentration unit) | Abbreviation | Compound category (severity) | Concentration level |  |  |  |  |  | Experiment batch |
| --- | --- | --- | --- | --- | --- | --- | --- | --- | --- |
|  |  |  | 1 | 2 | 3 | 4 | 5 | 6 |  |
| Amiodarone (μM) | AMI | DILI compound (8) | 0.8 | 4.0 | 8.1 | 16.2 | 32.3 | 64.6 | 1 |
| Diclofenac (μM) | DIC | DILI compound (8) | 10.1 | 50.5 | 101.0 | 202.0 | 404.0 | 606.0 | 1 |
| Ketoconazole (μM) | KET | DILI compound (8) | 3.3 | 6.6 | 33.0 | 65.9 | 131.9 | 263.8 | 1 |
| Nefazodone (μM) | NEF | DILI compound(8) | 1.9 | 3.9 | 19.7 | 39.5 | 79.0 | 100.0 | 1 |
| Nitrofurantoin (μM) | NIT | DILI compound (8) | 6.0 | 30.0 | 59.9 | 119.9 | 239.7 | 359.6 | 1 |
| Nitrofurantoin (μM) | NIT2 | DILI compound (8) | 6.0 | 30.0 | 59.9 | 119.9 | 239.7 | 359.6 | 2 |
| Phenytoin (μM) | PHE | DILI compound (8) | 10.9 | 54.3 | 108.7 | 217.3 | 869.2 | 1086.5 | 1 |
| Tolcapone (μM) | TOL | DILI compound (8) | 10.9 | 22.0 | 109.9 | 219.9 | 439.7 | 879.4 | 1 |
| Troglitazone (μM) | TGL | DILI compound (8) | 6.4 | 31.8 | 63.6 | 127.1 | 254.2 | 381.3 | 1 |
| Valproat (μM) | VPA | DILI compound (8) | 121.4 | 606.9 | 1213.8 | 2427.5 | 4855.1 | 7282.6 | 1 |
| Ciproflaxin (μM) | CIP | DILI compound (7) | 3.3 | 16.5 | 32.9 | 65.8 | 131.6 | 197.4 | 1 |
| Cyclosporin A (μM) | CYA | DILI compound (7) | 0.2 | 1.0 | 4.0 | 8.0 | 12.0 | 16.0 | 1 |
| Azathioprine (μM) | AZA | DILI compound (5) | 0.3 | 1.7 | 3.4 | 6.8 | 20.4 | 27.2 | 1 |
| Acetamonophen (μM) | ACE | DILI compound (5) | 69.5 | 347.4 | 694.7 | 1389.5 | 2778.9 | 4168.4 | 1 |
| Omeprazole(μM) | OMZ | DILI compound (4) | 4.7 | 23.5 | 47.0 | 94.0 | 188.0 | 282.1 | 1 |
| Cisplatin (μM) | CDDP | DILI compound (3) | 1.0 | 2.5 | 5.0 | 10.0 | 15.0 | 20.0 | 1 |
| Etoposide (μM) | ETO | DILI compound (3) | 1.0 | 5.0 | 25.0 | 50.0 |  |  | 2 |
| Famotidine (μM) | FAM | DILI compound (3) | 0.8 | 4.1 | 8.2 | 16.3 | 32.6 | 48.9 | 1 |
| Verapamil (μM) | VPL | DILI compound (3) | 0.5 | 2.6 | 5.1 | 10.2 | 30.5 | 50.9 | 1 |
| Buspirone (μM) | BUS | DILI compound (3) | 0.0 | 0.1 | 0.2 | 0.3 | 0.7 | 1.0 | 1 |
| Thapsigargin (μM) | THAP | ER stress inducer | 0.1 | 0.5 | 2.5 | 5.0 |  |  | 2 |
| Tunicamycin (μM) | TUN | ER stress inducer | 0.2 | 1.0 | 5.0 | 10.0 |  |  | 2 |
| CDDO (μM) | CDDO | Oxidative stress inducer | 0.0 | 0.1 | 0.5 | 1.0 |  |  | 2 |
| Diethyl maleate (μM) | DEM | Oxidative stress inducer | 3.0 | 16.0 | 80.0 | 160.0 |  |  | 2 |
| Sulfoephene (μM) | SUL | Oxidative stress inducer | 0.6 | 3.0 | 15.0 | 30.0 |  |  | 2 |
| Mitomycin C (μM) | MIT | DNA damage inducer | 1.2 | 6.0 | 30.0 | 60.0 |  |  | 2 |
| Rotenone (μM) | ROT | Mitochondria toxicant | 0.0 | 0.1 | 0.4 | 2.0 |  |  | 2 |
| IL-1β (ng/ml) | ILβ | Cytokines | 2.0 | 10.0 | 50.0 | 200.0 |  |  | 2 |
| TGFβ1 (ng/ml) | TGFβ | Cytokines | 0.5 | 2.0 | 10.0 | 40.0 |  |  | 2 |
| TNFα (ng/ml) | TNFα | Cytokines | 0.2 | 1.0 | 5.0 | 20.0 |  |  | 2 |
| Activin A (ng/ml) | ActA | growth factors | 0.2 | 1.0 | 5.0 | 20.0 |  |  | 2 |
| BMP4 (ng/ml) | BMP | growth factors | 0.2 | 1.0 | 5.0 | 20.0 |  |  | 2 |
| EGF (ng/ml) | EGF | growth factors | 0.2 | 1.0 | 5.0 | 20.0 |  |  | 2 |
| FGF1 (ng/ml) | FGF1 | growth factors | 0.5 | 2.5 | 12.5 | 50.0 |  |  | 2 |
| HGF (ng/ml) | HGF | growth factors | 0.5 | 2.0 | 10.0 | 40.0 |  |  | 2 |
| Serum (FBS) starvation (%) | Str | growth factors | 0.0 | 0.1 | 0.5 | 2.0 |  |  | 2 |
| Wnt3a (ng/ml) | Wnt | growth factors | 0.2 | 1.0 | 5.0 | 20.0 |  |  | 2 |
| Melatonin (μM) | MEL | Negative compound | 2.6 | 12.9 | 25.8 | 51.7 | 103.3 | 155.0 | 1 |
| N-Acetylcysteine (μM) | NAC | Negative compound | 1000.0 | 2000.0 | 4000.0 | 6000.0 | 8000.0 | 10000.0 | 1 |
| Vancomycin (μM) | VAN | Negative compound | 1.4 | 6.9 | 13.8 | 27.6 | 55.2 | 82.8 | 1 |
| Vitamin C (μM) | ASC | Negative compound | 70.0 | 349.8 | 699.5 | 1399.1 |  |  | 1 |

### 2.4 RNA sequencing and transcriptomic data analysis

The sequencing was performed deploying the human whole transcriptome library. The probe alignment for whole transcriptome gene set was performed by BioClavis. Briefly, FASTQ files were aligned using Bowtie, allowing for up to 2 mismatching in the target sequence. This pipeline applies several quality controls with mapped/unmapped reads, replicate clustering, and sample clustering^19^. Furthermore, we perform sample quality control steps to exclude samples with lower quality of the sequencing outcome (Suppl. figure 1). Only the correctly mapped counts (raw count values) were further processed. We first excluded samples with library size (the sum of raw count values) lower than 500,000 counts. Then, the raw count values were normalized with the CPM method. In addition, the replicate correlation for each sample towards the mean of the same conditions was evaluated and the samples showing lower than 0.95 Pearson correlation values were eliminated. The samples passed these QC steps were further used for log2 fold change calculation^20^. For the log2 fold change calculation, the normalized count values of each probe from the treated samples were compared to the values of the DMSO (except for cisplatin, N-acetyl cysteine, cytokines, growth factors, and solvents [DMSO, DMSO2, PBS, and PBSh] – compared to PBS [12%], PBSh [20%], and DMEM respectively) from the same time point. The log2 fold change values were then used as input for the WGCNA. Complementary, we defined the numbers of the differentially expressed genes using the thresholds: log2 fold change > |0.1| & adj-p value < 0.01. The transcriptomic data have been made available on https://www.ebi.ac.uk/fg/annotare/, with accession number E-MTAB-11555 for the first transcriptomic batch and E-MTAB-11578 for the second transcriptomic batch.

### 2.5 Gene network generation using the weighted gene co-regulated network analysis

Prior to applying the WGCNA approach to the processed data, the most significant probes based on the adjusted p-values were selected resulting in the one probe measurement for each gene. Furthermore, the goodSampleGenes^20^ function was applied to the data matrix to eliminate the gene with non-meaningful log2 fold change. Finally, to identify co-expressed genes from the PHH data, we used the WGCNA R package version 1.51^20^ and applied it to a matrix consisting of 260 rows (experimental conditions being a combination of compound-time-dose) and 11,153 columns (log2 fold change values for genes). We generated unsigned gene modules (enabling the clustering of co-induced and co-repressed genes), and selected the optimal soft power threshold maximizing both the scale-free network topology using standard power-law plotting tool in WGCNA. We selected 9 as the optimal soft-power parameter (Suppl. figure 3A). For each experiment, the eigengene score (EGs, or module score) was calculated which was derived from the log2 fold change values of their composing genes (Supp. table-tab 1). Briefly, this protocol consisted of performing PCA on the gene matrix of each module, normalizing the log2FC across the entire dataset using Z-score conversion, the 1st principal component corresponds to the EGs^14^. To facilitate the comparison between modules across treatment, the raw module score was normalized to unit variance (fraction between each module score and its standard deviation across the entire dataset). The EGs indicated the magnitude of activation or repression induced by a given treatment (Supp. table-tab 2). We further refined the modules built by merging similar modules (those having correlation of their EGs≥ 0.8), and obtained 288 modules (Suppl. figure 2B). Modules were annotated for their cellular responses based on the GO database by performing gene set enrichment using the enrichmentAnalysis function^21^ (Supp. table-tab 3). The most significant annotation based on the adjusted p-values was then chosen to represent the cellular responses of the modules. For every gene in a module, the correlation was calculated between the log2 fold change versus the eigengene score of its parent module across the 260 experiments (termed ‘corEG’). The gene with the highest correlation (so called ‘hub gene’) was the most representative of the entire module matrix and show stronger connection to the other gene memberships (Supp. table-tab 1). Preservation between the modules structure obtained with the HepG2 data set and the PHH TG-GATEs was performed using preservation statistics calculated with the WGCNA R package and thresholds for interpretation were adopted from the relevant literature^22^. Module showing Z summary >= 2 was considered moderately preserved, >= 10 highly preserved. A lower median rank indicates higher preservation.

### 2.6 Correlation analysis between module activation and cell death outcomes

The correlation analysis was applied to the module scores and the cell death outcomes from the previous study^17^. The cell death data was derived using live cell imaging capturing the fraction of the cell death indicated by with the propidium iodide (PI) staining for necrosis and annexin V (AnV) staining for apoptosis. The cell death data at 8, 24, 48, and 58 hours were selected for the correlation analysis. The data points with shared exposure conditions (the same compound and concentration) in both module scores and cell death outcome were selected. The correlation analysis was applied to each compound for every cell death type (necrosis and apoptosis) from the previously generated cell death data^17^ using Pearson correlation method in which the correlation value was composed of the outcomes from the shared concentration from both dataset. The positive correlated modules were then selected with the thresholds of correlation adjusted p-values < 0.1, correlation score > 0.5, EGs > 2 at least in one data point, and > 4 DILI compounds in which the correlation outcomes passing the thresholds. The genes inside the modules for further validation were selected based on the thresholds of adjusted p-val < 0.1 and log2 fold change > 2 at least in one condition.

### 2.7 RNA interference screen and live cell imaging for cell death determination

Among the high upregulated gene memberships of the high cell death-correlated modules, we selected the 67 target genes based on the availability of the siRNAs from the drugable genome library. siRNAs against targeted genes were purchased from Dharmacon (ThermoFisher Scientific) as siGENOME SMARTpool reagents, as well as in the form of individual siRNAs (Supp. table-tab 4). HepG2 cell suspensions were transiently transfected with the mixture of siRNAs (50 nM) and INTERFERin (Polyplus) in DMEM high glucose and seeded in Greiner black μ-clear 96 well plates, at 25,000 cells per well. The medium was refreshed 24 hours post-transfection and compound exposures were performed 48 hours afterward. siGENOME non-targeting pool #1 (siNo1) and mock (INTERFERin containing medium) condition were used as the control. On the day of the exposure, the cells were incubated with medium containing 100 ng/ml live Hoechst for 2 hours. The medium was then refreshed with the medium containing 0.2 μM propidium iodide (PI) and 1:2000 annexin V-Alexa 633 (AnV). Sequentially, the compounds were added to the plate to reach the desired concentration. For the RNAi experiment, we used nitrofurantoin (360 - 480 μM) and nefazodone (39.5 - 79 μM). The plates were imaged at 24, 48, and 72 hours after exposures using a Nikon TiE2000 confocal laser microscope (laser: 647 nm, 540 nm, and 408 nm), equipped with automated stage and perfect focus system. During the imaging, the plates were maintained in humidified atmosphere at 37 degrees Celsius and 5% CO_2_ /air mixture. The imaging was done with 20x magnification objective.

### 2.8 Image analysis

The images were manually sorted to exclude images which did not fulfill the criteria for analysis: non-biological background (fluorescent fibers), loss of nuclear signal, and out-of-focus images. The quantitative image analysis was performed with ImageJ version 1.52p and CellProfiler version 2.2.0. Firstly, the nuclei per image were segmented with watershed masked algorithm on ImageJ and thereafter processed with an in house developed CellProfiler module module^23,24^. The results were stored as HDF5 files. Data analysis, quality control, and graphics were performed using the in-house developed R package h5CellProfiler. The nuclear Hoechst33342 intensity levels, nuclear area, PI area, and Annexin V area were measured at the single cell level. The number of PI and AnV positive cells were determined based on the count of cells with higher than 10 % overlapping between nuclear area and PI/AnV area. For the RNAi experiments, the z-score of the cell death was calculated with this formula :

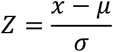

*x* = cell death value of each siRNA

*μ* = mean of the population (from the same compound, time point, and cell death type)

*σ* = standard deviation of the population

### 2.9 Data representation

The graphical representation of the results were generated or modified with Illustrator CS6 and R (ggplot2^25^ and pheatmap^26^). Correlation analysis was performed with the R function from the package Hmisc^27^. The hierarchical clustering is performed with “*Ward D2*” algorithm applied to the Euclidian distance between measured variables. Specifically for compound clustering based on the module activity, the clustering was subjected to cutree() function at the height of 20 to define the groups.

## Results

### Transcriptomic perturbations in the HepG2 cell line exhibit different response patterns between DILI and and non-DILI compounds

We performed an extensive toxicogenomics study in HepG2 cells exposed to different DILI compounds at different concentrations for varying time points (Table 1). We also included several cytokines and growth factors as well as defined reference compounds for toxicity relevant cellular stress response pathways to both broaden the diversity of xenobiotic-induced transcriptional responses and to provide benchmark information on important cellular response to stress. Several compounds were also added as negative controls that showed no cellular stress response modulation in HepG2 cells in previous studies.^17^ Samples were processed for high throughput transcriptomic (HTTr) based on targeted whole genome RNA sequencing using the TempO-Seq technology. TempO-Seq data showed overall sufficient library size across as well as replicate correlation for the different treatments (Suppl. Fig. 1). Differential expressed genes (DEGs) for each condition were determined with a defined threshold: adjusted-pval <0.01 and log2FC > |0.1| cutoff. The aggregated number of DEGs, based on the sum of the unique DEGs for all concentrations from each compound, reflected stronger transcriptional responses by the various DILI and reference compounds, but more limited responses for cytokines and growth factors (Figure. 1B). Comparing responses between two separate experiments (batches) with nitrofurantoin (NIT and NIT2) indicated that the inter-batch variation with respect to the number of DEGs and fold change was minimal (Figure. 1B and Suppl. Fig. 2A). Furthermore, most transcriptomic responses induced by the DILI compounds and positive compounds showed clear time dependence (Figure 1B) with strongest response observed at 24 hours. However, the transcriptomic responses of the cytokines and growth factors were more prominent at 8 hours (Suppl. Figure 2B – examples FGF1, HGF, and EGF). In addition, the transcriptomic perturbations exhibited a concentration-dependence with higher numbers of DEGs at increasing concentration of DILI compounds (Figure 1C – example for nitrofurantoin). Additionally, the principal component analysis plots showed different clusters of samples based on the compound categories, time points, and concentration level, indicating different transcriptomic profiles upon the exposure of the different compound categories at different concentration and time points (Figure 1D). Consistent with the number of DEGs, most of the DILI compounds and positive compounds at the higher dose (dose level >= 4) exhibited clear separation from negative compounds (24 hours PCA). The responses of the cytokines and growth factors clustered separately from other compounds at 8 hours but not at 24 hours, confirming the early transcriptomic perturbation. These findings suggest that each compound category induces distinctive transcriptomic responses. Overall, the gene-level data suggested that the compound sets used induced robust gene expression responses with considerable diversity amongst treatments.

### Cellular stress response-related gene networks show coherent modulation upon the exposure of positive control compounds

Previous work has shown that co-expression analysis can yield expected and novel insights into biological responses to cellular stress^,14,15^. In order to identify gene networks associated with cellular responses, we performed weighted gene co-regulated network analysis (WGCNA) on the transcriptomic data to identify co-regulated sets of genes (modules). We obtained 288 modules compose of 14,359 genes (Suppl. figure 3B and Supp. table-tab 1) which also resulted in 98% reduction of the dimensionality of the gene expression originally derived from the readout of ∼21,111 probes. Moreover, the WGCNA also reduced the inter-batch variation as indicated by the higher correlation of the transcriptional changes upon nitrofurantoin treatment at the module level (Pearson R : 0.94) compared to the gene level (Pearson R : 0.74). The batch correlation at the module level was comparable to the correlation at the DEGs level (Pearson R : 0.96) (Suppl. figure 2A). This indicated that the module network derived from WGCNA was able to reduce dimensionality and eliminate the noise effect due to non-perturbed and/or low expressed genes.

Since our goal was to identify causal links among genes, biological responses and liver injury using HepG2 as an in vitro model, we first wanted validate the approach by determining if expected stress responses for model compounds were reflected in gene networks and their biological annotations. We focused on the activity of the modules that were annotated for biological response know to be associated with DILI-related cellular stress responses^4,28^: Module annotations were based on the gene ontology enrichment results for each individual module (Supp. table-tab 3) and supported by the presence of well-established gene memberships in the adaptive stress responses. Using this approach, we selected 4 modules for further analysis: HepG2:75-inflammation, HepG2:46-oxidative stress, HepG2:33-DNA damage, and HepG2:38-ATF4-CHOP complex (ER stress-related). The latter module is a generalized stress response target by the EiF2-alpha kinases^29^ often associated via CHOP/DDIT3 as a downstream effector arm of ER stress^30^. As expected, genes belonging to the same modules showed similar regulation patterns indicative of co-regulation while responses between genes from different modules varied (Figure 2A). This effect was particularly appreciable upon the exposure to the reference compound: TNFα for HepG2:75-inflammation, DEM for HepG2:46-oxidative stress, mitomycin C for HepG2:33-DNA damage, and tunicamycin for HepG2:38-ER stress (Figure 2A). Thus, individual HepG2 module genes respond in a coordinated fashion to compounds expected to activate the biological processes derived empirically solely from the module enrichment annotations.

**Figure 2.**
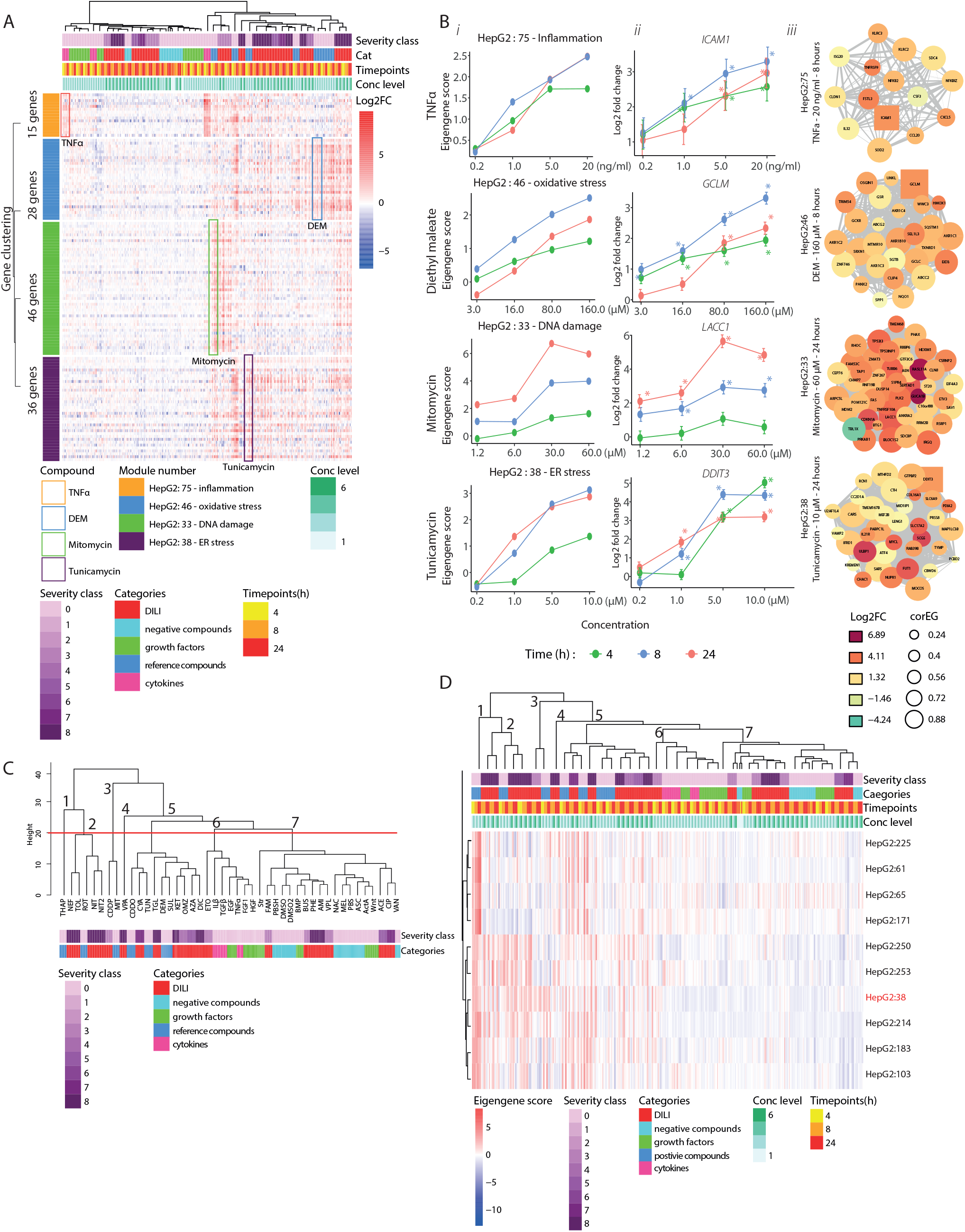
Gene network activation of WGCNA-based modules by DILI compounds. (A) A heatmap (n=3) exhibiting the modulation of the gene memberships of WGCNA:HepG2:75, WGCNA:HepG2:46, WGCNA:HepG2:33, WGCNA:HepG2:38. The highlighted parts (colored boxes) show the modulation of the gene upon the positive control exposure : orange – TNFα, blue – DEM, green – mitomycin, purple – tunicamycin. The heatmap contains 4 variables on the columns indicated by distinctive color groups: severity class of the compound, compound categories, time points, and dose level. These variables describe the exposure condition applied to the cells. Each row of the heatmap shows the log2 fold change values of each module memberships from every sample where red color shows upregulation and blue color shows downregulation. The color on the row indicates the modules (orange: HepG2:75, blue: HepG2:46, green: WGCNA:HepG2:33, and purple: HepG2:38). The clustering of the heatmap is performed using the “*Ward d2*” algorithm applied to the Euclidian distance between aggregated variables (column clustering: mean of the log2 fold change values of the dose levels and time points per compound, row clustering : mean of the log2 fold change values of the genes per module). (B) The dose response plots of stress response related modules upon the exposure of the specific reference compounds: HepG2:75 – inflammation (TNFα), HepG2:46 - oxidative stress (DEM), HepG2:33 - DNA damage (etoposide), HepG2:38 – ER stress (tunicamycin) (i). The dose response plots of the hub gene of each stress response related module upon the exposure of the specific reference compounds: HepG2:75 – *ICAM1*, HepG2:46 - *GCLM*, HepG2:33 - *TNFRS10A*, HepG2:38 - *MTFD2*. The color of the plot represent the time points, the error bars indicate standard error mean (SEM) values, n=3. Stars indicate the significant upregulation with adjusted p-value < 0.01 (ii). The overview of the gene memberships’ activities of the stress response modules upon the exposure of the positive compound at the particular time point and concentration. The box node indicate the hub gene (the gene with the highest correlation to the parent module). The color of the nodes represents the modulation and the size of the nodes represent the module correlation (This figure is enlarged in Suppl. figure 3C) (iii). (C) The compound clustering based on aggregated mean eigengene scores for every time point and dose level per compound. The hierarchical clustering is performed using “*Ward D2*” algorithm applied to the Euclidian distant between aggregated values. The compounds are annotated with the severity and categories showed by the color bars. The determination of the clusters was set at the height 20 with the cuttree() function. (D) A heatmap showing the cluster of the modules annotated as the endoplasmic reticulum activities and ER stress responses. The heatmap contains 4 variables indicated by distinctive color groups : severity class of the compound, compound categories, time points, and dose level. These variables describe the exposure condition applied to the cells. Each row of the heatmap shows the eigengene values where red color shows activated and blue color shows deactivation. The clustering of the columns is performed with the same manner as compound clustering (C). HepG2:38-ATF-CHOP ER stress is highlighted with red.

Having established that individual module genes respond as expected, we next asked whether the module eigengenes score (EGs), which is a single value reflecting the behavior or all module genes, responded in a similar manner using the four reference compounds. We compared the EGS to the fold change for the module hub gene, (the gene with the highest module correlation or Core EGS; Supp. table-tab 1). The stress response module EGS showed time- and concentration-dependent responses; module activity was typically already high at 8 hours, except for HepG2:33 which showed strongest induction at 24 hours. As expected, the upregulation of the hub genes for each of these four modules showed a similar concentration and temporal response pattern to their parent modules (Figure 2Bii). The modulation of the gene members for each stress response related module was illustrated in the arrangement of nodular networks with the square node as the hub gene (Figure 2Biii-enlarged in Suppl. figure 3C). Yet, *DDIT3*, the HepG2:38 hub gene, showed strong upregulation already at 4 hours with a peak at 8 hours. However, we observed that >50% of the HepG2:38 gene members showed stronger upregulation at 24 hours significantly contributing to the linear time-dependency of the parent module (Suppl. figure 4A). Based on these findings, we conclude that the modulation of the different cellular stress responses, based on the module enrichment, is reflected by the module EGS and the member genes exhibited the expected coherent activation patterns after treatment.

### Co-expression modules capture compound class differences and highlight candidate cellular mechanisms underlying high DILI risk compounds

We further interrogated the activity of the modules beyond the stress response-related modules in order to uncover mechanisms in HepG2 cells activated after DILI compound exposure. Hierarchical cluster analysis revealed that both compounds and modules clustered according to the EGs activity (Suppl. figure 4B). There were 7 compound clusters (Figure 2C). Clusters 1-5 contained the reference compounds and DILI compounds with high DILI severity class. Cluster 6 consisted of the cytokines and growth factors. Clusters 7 mainly included the low severity class DILI compounds as well as DILI negative compounds. Overall, compound clusters 1-5 showed stronger (de)activation of the various modules compared to clusters 6-8 (Suppl. figure 4B). Of note, acetaminophen was grouped in cluster 8 and showed minor module activation, which was anticipated since HepG2 cells have little cytochromeP450 2E1^31,32^ required for the formation of the toxic metabolite. As an example for the module clustering, we highlighted one module cluster (see red box in Suppl. figure 4B and detailed in Figure 2D). Interestingly, this cluster consisted of the modules annotated for the endoplasmic reticulum related responses or/and stress (including HepG2:38-red, HepG2:61, HepG2:225, HepG2:65, HepG2:171); modules in this clusters were strongly activated by high severity DILI compounds (clusters 1-4) including nefazodone, tolcapone, and nitrofurantoin (Figure 2D).

We then identified the most activated and repressed modules (Table 2) across the entire compound set to investigate the strongest modulated cellular mechanisms upon DILI compound treatment. In order to select strongly perturbed modules, the median scores of the EGs across the conditions were calculated. We further counted the number of conditions that perturbed the modules resulting in EGS > 2 for activation and EGS < -2 for repression. Based on these two criteria, the modules were sorted and top 20 activated and repressed modules were selected. Interestingly, all of the stress response-related modules were among the top 20 activated modules along with other ER stress related (HepG2:38) and inflammation related responses (HepG2:140). Moreover, cytoskeletal re-organization (HepG2:101, HepG2:158, HepG2:196) and small molecule activity (HepG2:206, HepG2:235) were also highly activated. On the other hand, the top 20 top repressed modules consisted of the modules annotated for organelle biogenesis (HepG2:24, HepG2:37, HepG2:71), mitochondria (HepG2:70, HepG2:282), metabolism activity (HepG2:58, HepG2:251), and cell cycle activation (HepG2:10, HepG2:252). Altogether, these co-expression modules could highlight the cellular processes that are most often involved in the hepatocellular mechanisms of DILI.

**Table 2.** Top 20 most modulated gene networks upon the exposures of the tested compounds. The top 20 most repressed and activated upon exposure of the tested compounds. The selection was based on the calculation of the median scores of the EGS across all conditions and the number of conditions (count) which perturb the modules resulting in EGS > 2 for activation and EGS < -2 for repression. Based on these two criteria, the modules were sorted and top 20 activated and repressed modules were selected

| Module | Modulation | Annotation | Hub Gene | Preservation in PHH |
| --- | --- | --- | --- | --- |
| HepG2:1 | De-Activated | G-protein-coupled receptor activity, transmembrane signaling receptor activity, signaling receptor activity | *KRTAP16-1* | Non-preserved |
| HepG2:4 | De-Activated | polyketide metabolic process | *GJB1* | Preserved |
| HepG2:10 | De-Activated | cell cycle process, DNA replication, DNA metabolic process, mitotic cell cycle process | *SAC3D1* | Preserved |
| HepG2:24 | De-Activated | nucleosome assembly, chromatin assembly | *HIST1H2AD* | Preserved |
| HepG2:32 | De-Activated | positive regulation of glycogen biosynthetic process, leukocyte activation involved in inflammatory response | *ZNF607* | Preserved |
| HepG2:36 | De-Activated | rough endoplasmic reticulum lumen, DNA binding | *ZNF594* | Preserved |
| HepG2:37 | De-Activated | positive regulation of mitochondrial membrane potential | *MSS51* | Non-preserved |
| HepG2:58 | De-Activated | cholesterol biosynthetic process | *MVD* | Preserved |
| HepG2:69 | De-Activated | thyroid stimulating hormone | *FOXD4L3* | Non-preserved |
| HepG2:70 | De-Activated | mitochondrial membrane part, integral component of organelle membrane | *COA3* | Preserved |
| HepG2:71 | De-Activated | endoplasmic reticulum lumen, post-translational protein modification, histamine metabolic process | *SERPINA10* | Preserved |
| HepG2:86 | De-Activated | protein localization to plasma membrane, COPII adaptor activity | *TEX261* | Non-preserved |
| HepG2:123 | De-Activated | procollagen-proline 4-dioxygenase activity, oxidoreductase activity | *EFCAB3* | Non-preserved |
| HepG2:137 | De-Activated | positive regulation of cellular response to hypoxia, hypoxia-inducible factor-1alpha signaling pathway | *RNF170* | Non-preserved |
| HepG2:147 | De-Activated | endoplasmic reticulum-Golgi intermediate compartment, endoplasmic reticulum membrane, MHC class I protein complex assembly | *HINT2* | Preserved |
| HepG2:150 | De-Activated | xylulokinase activity, IkappaB kinase complex binding | *LARS2* | Non-preserved |
| HepG2:175 | De-Activated | RNA methyltransferase activity, cellular response to peptide, DNA-dependent ATPase activity | *METTL3* | Non-preserved |
| HepG2:251 | De-Activated | UDP catabolic process, pyrimidine nucleoside diphosphate catabolic process | *JRK* | Non-preserved |
| HepG2:252 | De-Activated | positive regulation of cell cycle, nuclear division, organelle fision, cell division | *CCNF* | Preserved |
| HepG2:282 | De-Activated | mitochondrial membrane fusion, desmosome assembly, positive regulation of mitochondrial translation | *RCC1L* | Non-preserved |
| HepG2:3 | Activated | regulation of signaling, regulation of cell communication, regulation of response to stimulus, regulation of signal transduction | *UBQLN1* | Preserved |
| HepG2:33 | Activated | signal transduction by p53 class mediator, apoptotic process, cellular response to UV | *TNFRSF10A* | Preserved |
| HepG2:38 | Activated | CHOP-ATF4 activity, ER stress response, ER stress related apoptotsis | *MTHFD2* | Preserved |
| HepG2:43 | Activated | negative regulation of blood vessel remodeling | *PALM2* | Non-preserved |
| HepG2:46 | Activated | oxidative stress, polyketide metabolic process, alditol:NADP+ 1-oxidoreductase activity | *GCLM* | Preserved |
| HepG2:52 | Activated | DNA-binding transcription repressor activity, RNA polymerase II specific, regulation of cellular metabolic process | *IBTK* | Preserved |
| HepG2:61 | Activated | response to endoplasmic reticulum stress, endoplasmic reticulum unfolded protein response, IRE1-mediated unfolded protein response | *DNAJB9* | Preserved |
| HepG2:75 | Activated | response to cytokine | *ICAM1* | Preserved |
| HepG2:83 | Activated | synapse assembly, positive regulation of transcription, DNA-templated | *PIEZO1* | Preserved |
| HepG2:84 | Activated | tubulin-tyrosine ligase activity, C-terminal protein-tyrosinylation, histidine-tRNA ligase activity | *SLC25A51* | Non-preserved |
| HepG2:101 | Activated | piccolo NuA4 histone acetyltransferase complex, nuclear part, prenyltransferase activity | *FOXQ1* | Non-preserved |
| HepG2:112 | Activated | KDEL sequece binding, autophagosome, TGFb receptor signalling activity | *PIP4K2C* | Preserved |
| HepG2:140 | Activated | notochord cell differentiation | *NDE1* | Non-preserved |
| HepG2:158 | Activated | cadherin binding, cell adhesion molecule binding | *SNHG8* | Preserved |
| HepG2:196 | Activated | actin cytoskeleton, homotypic cell-cell adhesion | *TAGLN* | Preserved |
| HepG2:206 | Activated | multicellular organism aging, aspartate-glutamate transport | *CREBRF* | Preserved |
| HepG2:214 | Activated | Atg1/ULK1 complex activation, ER stress, Macroautophagy | *SESN2* | Preserved |
| HepG2:228 | Activated | mitochondrion localization, neurotrophin production, negative regulation of PPAR signalling pathways | *REM2* | Non-preserved |
| HepG2:235 | Activated | adenylate-cyclase inhibiting adrenergic receptor activity, glycine transport | *CALCOCO2* | Non-preserved |
| HepG2:253 | Activated | dodecenoyl-CoA delta-isomerase activity | *ERRFI1* | Preserved |

### Stress response-related modules are preserved in primary human hepatocytes

Having established that modules reduce the complexity of gene expression data to interpretable mechanistically relevant biological responses, we want to determine if HepG2 modules were preserved in PHH a gold standard for human liver *in vitro* test systems.. We performed a preservation analysis of the HepG2 modules using as reference the TG-GATEs PHH modules as a reference^15^ (Figure 3). Based on the z-summary preservation scores, 87 modules (∼30%) of the HepG2 modules showed moderate (z-summary 2-10) to high preservation (z-summary >10) in the PHH dataset (Figure 3B). The top 10 most preserved HepG2 modules in PHH were annotated with multiple essential cellular processes and functions such as cell cycle and division (HepG2:12, HepG2:10, HepG2:8), signal transduction (HepG2:3), cellular biosynthesis and respiration (HepG2:58, HepG2:4, HepG2:17), and endoplasmic reticulum-related processes (HepG2:38, HepG2:65, HepG2:61). In addition to these modules, the modules related to cellular stress responses^23,31,33^ such as oxidative stress (HepG2:46), DNA damage (HepG2:33), inflammation (HepG2:75), and ATF-4-CHOP complex (HepG2:38) were also preserved in PHH (marked in red). This suggested that the expression of the genes of the aforementioned stress responses modules are also co-regulated in HepG2 and PHH.

**Figure 3.**
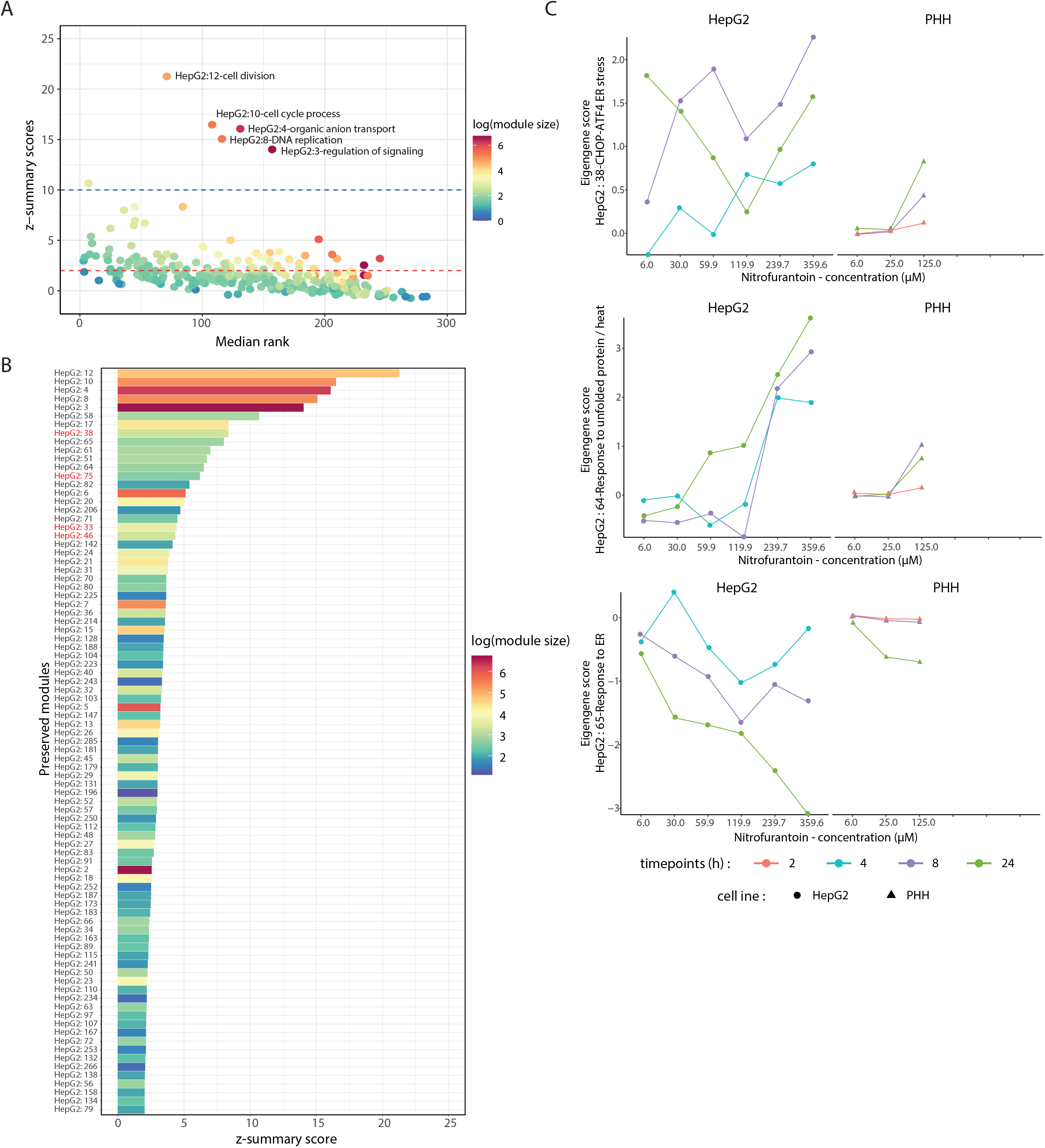
HepG2 module preservation analysis in the primary human hepatocytes (PHH). (A) The correlation plot between two module preservation parameters : z-summary scores and median rank. The red line in the y-axis intercept corresponds to z-summary score = 2 and the blue line to z-summary score = 10. Each dot represents each module in the plot. The color of the plots represents the module size in log scale. (B) The list of the HepG2 modules that are preserved in the PHH systems. The module written in red are the stress responses related modules. The color of the bar plots represents the module size in log scale. (C) Response of HepG2 and PHH upon nitrofurantoin exposure based on the HepG2:38, HepG2:64, and HepG2:65. The colors of the plot represent time points and the shapes of the dots indicate the cell lines.

To examine the similarity of the dynamics of preserved responses across the two test systems, we interrogate the dynamics of ER stress-related responses as an example exemplified with nitrofurantoin treatment (Figure 3C). In order to calculate the different module EGs for PHH, we used as input the log2 fold change derived from the responses of PHH as previously reported^34^. Module activation was observed for HepG2:38 and HepG2:64 (responses to unfolded protein and heat); module repression was observed for of HepG2:65 (response to ER stress). The temporal responses across test systems showed strong resemblance (Figure 3C). Yet, modules in HepG2 showed higher absolute EGs values compared to PHH; this was likely due to the higher log2 fold change values in HepG2 compared to PHH in relation to the higher compound concentration and the broader dynamic range of RNA sequencing compared to microarray^35^ (Suppl. figure 5A). Overall, for nitrofurantoin exposure, the module responses between HepG2 and PHH exhibited moderate similarity with 0.56 Pearson correlation score when the cells were exposed with comparable conditions (Suppl. figure 5B). As ER stress showed strong preservation, we evaluated this response based on the three preserved modules on another ER-stress inducing DILI compound, cyclosporine A. We observed similar direction of module activation between HepG2 and PHH (Suppl. figure 5C). Interestingly, for cyclosporine A, we found in both cells the activation of HepG:65 while HepG2:64 was not induced. In conclusion, we identified the gene networks that are preserved between HepG2 and PHH, and delineated the relevance of HepG2 observations for PHH.

### Modules highly correlated to cell death reveal candidate hepatocellular responses networks

. Ultimately, we wanted to establish direct causality between gene responding to cellular injury and cell death. As a first step, we focused on identifying module that showed a good correlation with cell death to assemble a list of candidate modules and genes. We identified modules that were linked to hepatocellular death by correlating the EGs of all modules with the live cell imaging of necrotic and apoptotic cell death readouts, propidium iodide and AnnexinV-Alexa633, respectively^17^. We performed the correlation analysis between modules’ EGs and the cell death outcomes measured at 8, 24, 48, and 58 hours, across all the compounds tested in order to identify common responses activated in the HepG2 in relation to cell death. The correlation between early module activities and cell death at 58 hours resulted in the highest numbers of significant correlation scores (adjusted p-values < 0.1; Suppl. figure 6A). As an example, nefazodone treatment caused most extensive cell death as well as strong transcriptomic perturbations; we observed high correlation between EGS of HepG2:38 and cell death markers at later time points (Figure 4A).

**Figure 4.**
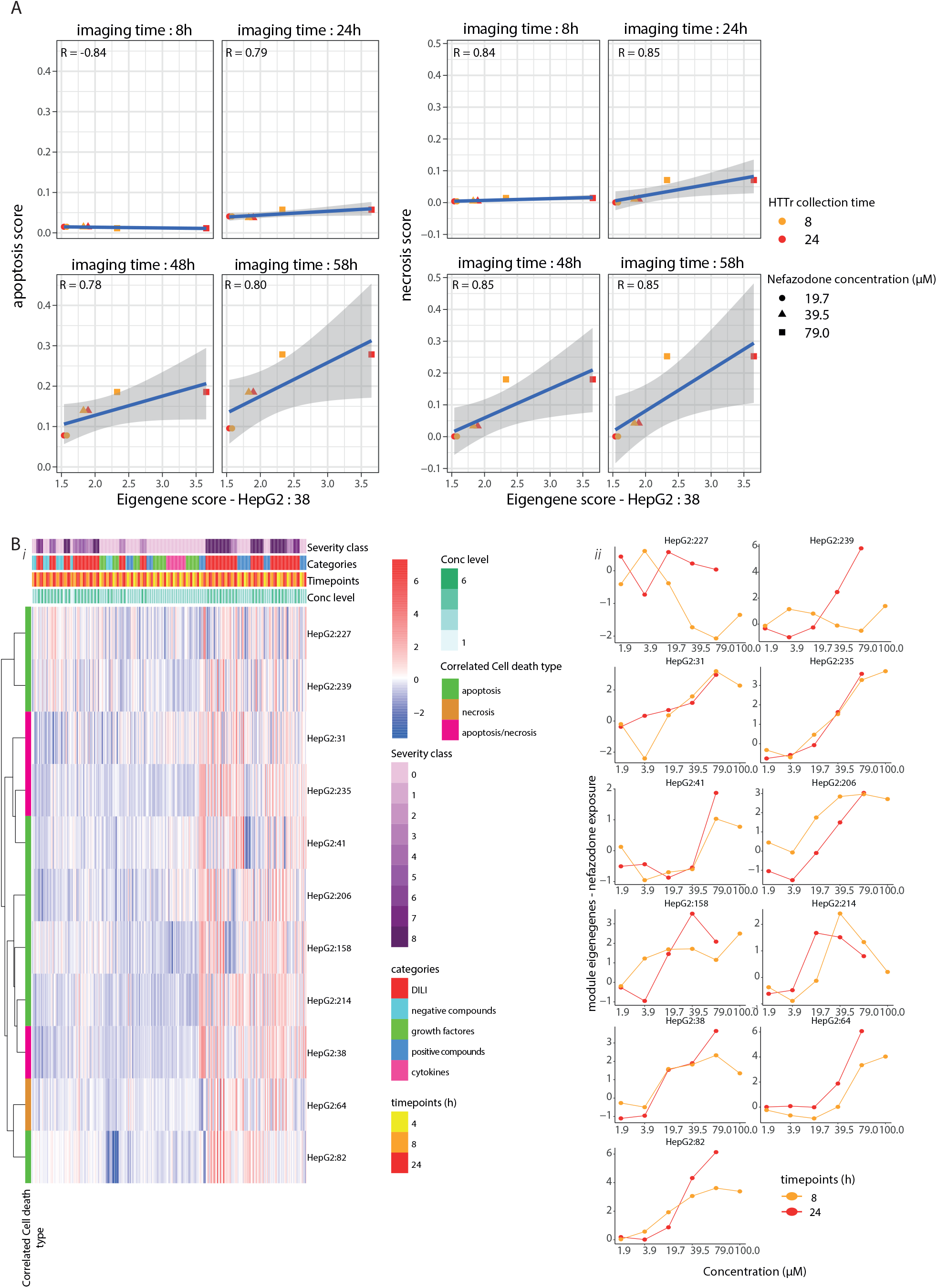
Module correlation analysis with cell death. (A) The correlation plots between eigengene score of HepG2:38 and percentage apoptosis (left) and necrosis (right) at designated cell death measurement time points (8, 24, 48, and 58 hours) on the cells exposed to nefazodone. The color of the dots correspond to the transcriptomic time point and the shape of the dots to the concentration. (B) A heatmap showing the eigengene scores of high correlated module to the cell death upon the exposure of the compounds. The heatmap contains 4 variables in indicated by distinctive color groups : severity class of the compound, compound categories, time points, and dose level. These variables describe the exposure condition applied to the cells. Each row of the heatmap shows the eigengene values of each module from every sample where each column of the heatmap indicates the sample bearing the identities based on the assigned variables. The modules are clustered using “*Ward D2*” algorithm. The color identity of the module indicates the correlated cell death type. The color of the heatmap representing the modulation of the gene networks (module) where red color shows activated and blue color shows deactivation (i). Dose response plots of the high cell death correlated-modules upon the exposure of nefazodone at 8 (orange) and 24 (red) hours (ii).

Since cell death onset was in particular observed at late time points, we further focused on the correlation analysis between the module activities (at 4, 8, and 24 hours) and cell death measured at 58 hours. We identified modules correlation scores with cell death outcomes for several DILI compounds (Supp. table-tab 5). Eleven modules passed the criteria of correlation adjusted p-values < 0.1, correlation score > 0.5, EGs > 2 at least in one data point, and > 4 DILI compounds in which the correlation outcomes passed these thresholds (Table 3). Ten of the high cell death modules were correlated with apoptotic and necrotic cell death and interestingly only one, HepG2:64, was significantly correlated with necrotic cell death. The activities of these modules showed prominent modulation mostly upon the exposure with severe DILI compounds (Figure 4Bi). We observed clear concentration response activation of these modules for the severe DILI compounds (example : Figure 4Bii-nefazodone and Suppl. figure 6B-nitrofurrantoin, troglitazone, and diclofenac). Since most modules associated with cell death were already activated at very early time points, this suggests that the activation of these modules may also be causally associated with cell death onset in HepG2 cells. The annotation of these modules showed broader cellular responses, some with known involvement in liver injury: ER stress related responses^33^ (HepG2:38, HepG2:64) and cytoskeletal re-organisation^36,37^ (HepG2:39, HepG2:158). In addition, cellular processes related to adenylate-cyclase (HepG2:235) were also highly correlated with cell death. Altogether, we identified key modules that exhibited high correlation with the occurrence of cell death.

**Table 3.**
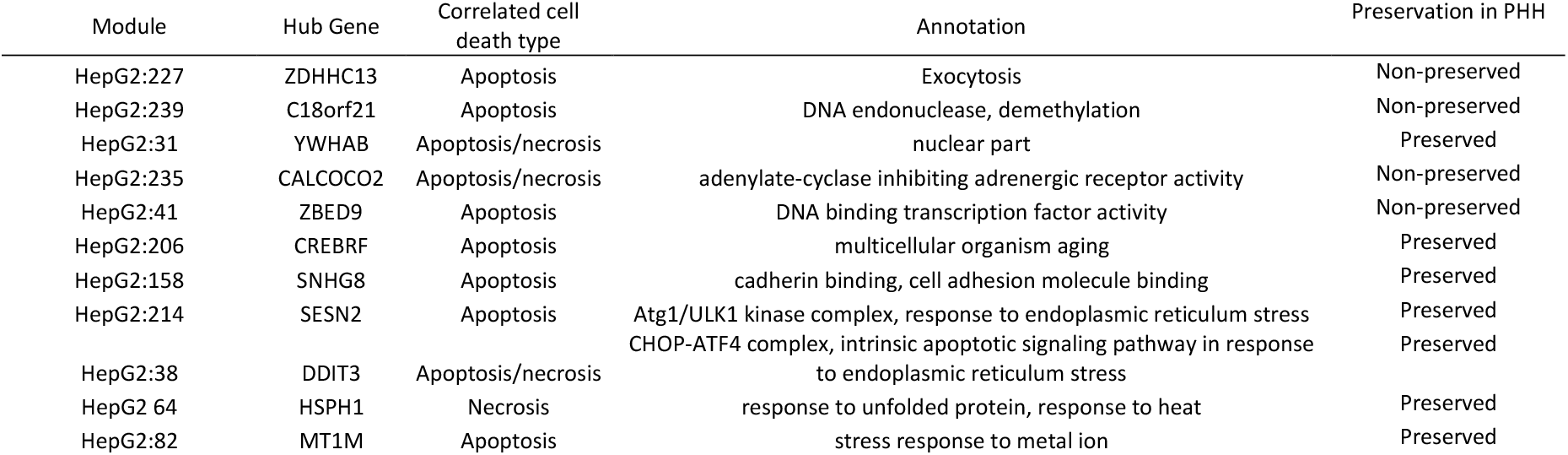
Modules with high correlation with cell death. These modules pass the threshold of correlation adjusted p-values < 0.1, correlation score > 0.5, EGs > 2 at lease in one data point, and > 4 DILI compounds in which the correlation outcomes passing the thresholds. The order is based on the clustering on Figure 4Bi.

### A decrease in cell death is found upon the perturbation of the gene memberships of the high cell death correlated modules

We reasoned that the module associations with cell death could be indicative for causal involvement. Therefore, the causality of the high correlated modules was further evaluated experimentally using RNA interference-based gene silencing of the gene members of the high correlated modules that showed strongest perturbations by DILI compounds (log2 fold change > 2 at least in one condition and p-adj < 0.1). As a proof-of-concept, from in total 141 genes (Supp. table-tab 6), we selected 67 gene targets to be perturbed based on availability in a drugable siRNA library, hence involved in cell signaling programs (Supp. table-tab 7). After siRNA knock down, HepG2 cells were exposed to nitrofurantoin and nefazodone, since these two DILI compounds showed strongest cell death onset and transcriptomic perturbation. Since *DDIT3* was also one of the 67 selected genes (Supp. table-tab 6), we used our HepG2-CHOP-GFP^23^ to capture the overall knock down efficiency within the experiment. Overall we anticipated that genes involved in protective adaptive responses would enhance cell death upon knock down, while genes that stimulate the onset of cell death would be protective upon knock down. The z-score analysis showed that knock down of some candidate genes reduced both apoptotic and necrotic cell death at 24 hours after the exposure with nefadozone and nitrofurantoin, while knock down of other genes enhanced cell death (Supp. table-tab 8). The candidate genes clustered into a cytotoxicity protective group after knock down, including *GTPBP2, HSPA1B, TSC22D3, SIRT1, IRF1*, and another 34 genes in the blue box as well as a cytotoxicity enhancing group including amongst others *DDIT3, MARCH6, SLC6A9, SLU7, HBP1, MYCL1* in the red box (Figure 5B). Interestingly, we found that the knock down of *DDIT3* known as the pro-apoptotic protein^38^ increased the cell death values upon the cell injury. Previous studies also reported the increase of the cell death upon the perturbation of DDIT3 expression^17,30^ suggesting different roles of this protein during cellular stress. We further focused on the target genes that reduced the cell death upon knock down for which cells persistently showed the decrease of cell death onset up to 72 hours of compound treatment. Cytotoxicity by nitrofurantoin was likely protected by GTPBP2 for prolonged time period. Nefazodone-induced cell death was most strongly protected by siGFPBP2, siIRF1 and siSIRT1 up to 72 hours. In conclusion, we discovered genes that are part of modules associated with cell death onset that are causality related to cell death onset or adaptive response, thus supporting the relevance of the statistical module activity associations with cytotoxic outcomes. Finally, we wanted to ensure that our selected genes were also modulated in PHH exposed to DILI compounds and for this used the TG-GATEs dataset^15^. In TG-GATEs dataset we identified 37 out of the 39 genes (HSP90AA1 and NGFR are not included in the PHH data) whose perturbation reduced cell death and evaluated their expression in PHH after DILI compound treatment (Suppl. figure 7). Five genes that showed most significant protection against cytotoxicity after siRNA treatment, - *GTPBP2*-HepG2:38, *HSPA1B*-HepG2:64, *IRF1*-HepG2:41, *SIRT1*-HepG2:31, and *TSC22D3*-HepG2:235 - exhibited prominent upregulation upon exposure with DILI compounds in PHH, in particular at the highest concentration at 24 hours (Figure 5D).

**Figure 5.**
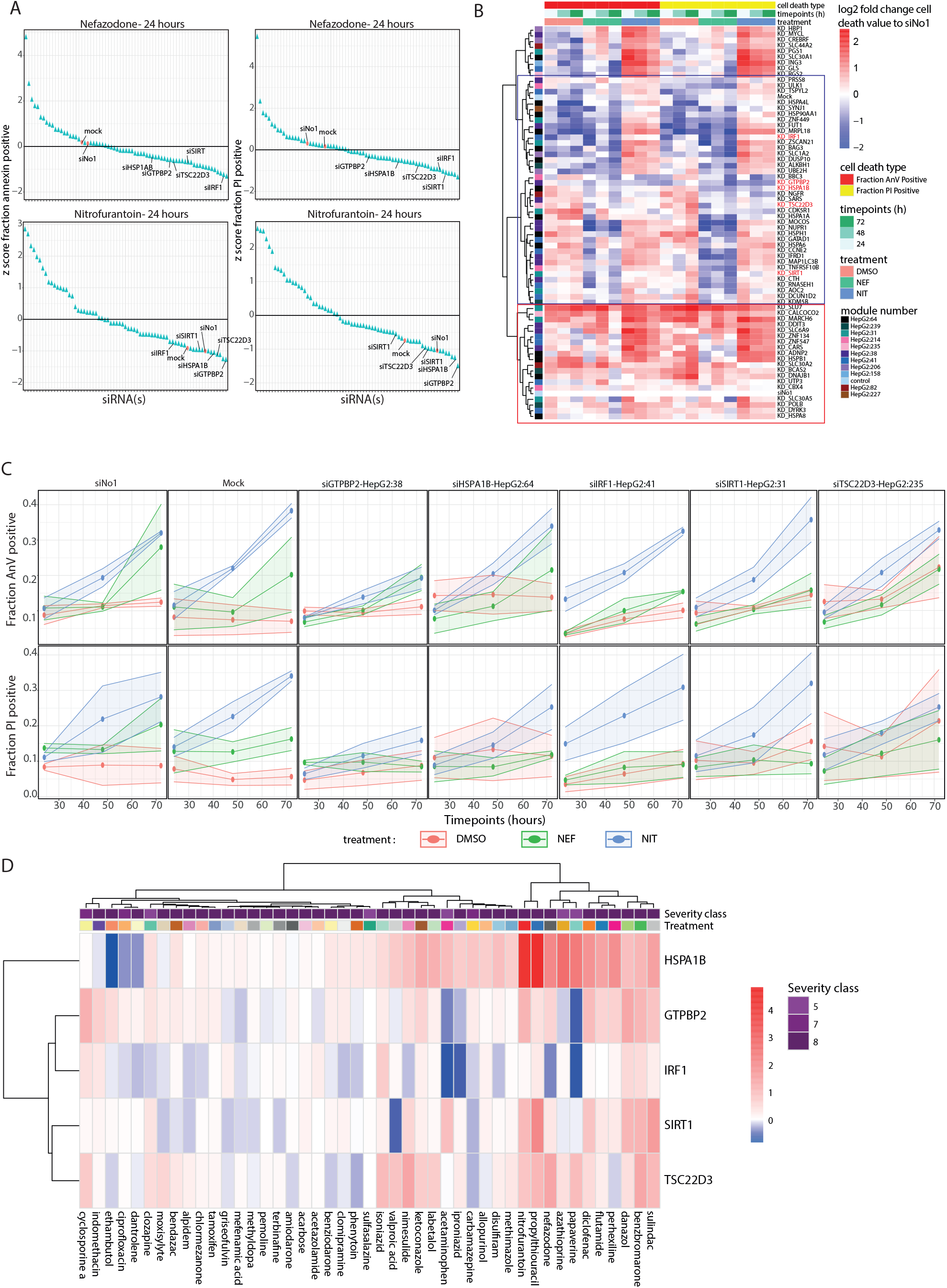
RNA interference to infer causal relationship of module gene memberships with cell death. (A) The dot plots showing the z-score of the cell death modulation in each perturbation compared upon nefazodone (top) and nitrofurantoin (bottom) exposure at 24 hours (left – annexin-V-Alexa633 readout, bottom – PI readout). The colors of the dots represent the type of the perturbation (red : control (mock [medium + INTERFERin] and siNo1 [medium + INTERFERin + scrambled siRNA), blue: siRNA targeting genes). (B) A heatmap (n=3) showing the log2 fold change cell death values compared to siNo1 of the HepG2 cells upon siRNA perturbation reducing the expression of the (highly upregulated) genes memberships of the high cell death-correlated modules. The heatmap consists of three legends: cell death type (red: fraction annexin-V positive cells; yellow: fraction PI positive cells), imaging time (green intensity: 24, 48 and 72 hours), and treatment (orange: DMSO; green: nefazodone (39.5 μM); blue: nitrofurantoin (360 μM)). The color on the row indicate the parent modules of the perturbed genes. The red intensity of the heatmap positively represents the cell death magnitude. The blue box contains the siRNAs that reduce the cell death and the red box contains the siRNA that increase the cell death. The red siRNAs highlight the highest cell death reduction upon the perturbation of the target genes. (C) The detailed overview of the perturbation of the hit genes (in red) resulting in the highest reduction of the cell death (top : fraction annexin-V positive cells, bottom: fraction PI positive cells). The color of the plot indicate the treatment (red: DMSO, green: nefazodone, blue: nitrofurantoin). The shadow of the plot representing the standard error mean (SEM), n=3. (D) The heatmap shows the expression in PHH of the 5 genes whose perturbation resulted in the highest reduction of the cell death fraction in HepG2 cells upon the exposure of nitrofurantoin (360 μM) and nefazodone (39.5 μM). The heatmap contains 3 variables on the columns indicated by distinctive color groups: severity class of the compound, time points, and dose level. These variables describe the exposure condition applied to the cells. Each row of the heatmap shows the log2 fold change values of each gene from every sample where red color shows upregulation and blue color shows downregulation. The column clustering of the heatmap is performed using the “*Ward d2*” algorithm applied to the Euclidian distance between aggregated variables (mean of the log2 fold change values from all the dose levels and time points per compound). The row clustering of the heatmap is performed using the “*Ward d2*” algorithm applied to the Euclidian distance between genes.

## Discussion

In this current study, we applied network-based analysis on a large novel concentration and time response transcriptomic dataset of HepG2 cells exposed to more than forty substances including 18 DILI compounds, 3 cytokines, 7 growth factors, 7 cellular stress responses pathway reference compounds, and 6 non-DILI compounds. Based on this transcriptomics dataset, we successfully established HepG2 cell-specific WGCNA-based gene networks encompassing the spectrum of cellular biological responses and defined the preservation of gene regulation between HepG2 and PHH. Key gene networks were identified that are associated with onset of cell death and candidate representatives from these networks were involved in modulation of cell death. These genes were also modulated by various DILI compounds in PHH, with most considerable changes observed for *HSPA1B, TSC22D3, GTPBP2* and *SIRT1*.

We established HepG2 gene networks to facilitate the interpretation of toxicogenomics datasets by reducing the data dimensionality, the ability to assign cellular function and responses for each network, and quantifying the biological activation of the gene networks. Our module-based approach captured the gene networks related to various cellular responses linked to the mechanisms of DILI such as stress responses^17,33^ (HepG2:46, HepG2:33), ER stress^39^ (HepG2:38, HepG2:64, HepG2:65), mitochondrial damage^40^ (HepG2:70, HepG2:282), and inflammation^41^ (HepG2:75, HepG2:140). Complementary, we identified the repressed cellular responses such as organelle biogenesis (HepG2 : 24, HepG2 : 37), small molecule metabolism activity (HepG2 : 58, HepG2 : 69, HepG2 : 251), and cell cycle activation (HepG2 : 10, HepG2 : 252) known to play a role in liver regeneration^42,43,44^. Importantly, preservation analysis showed that 30% of HepG2 modules are preserved in PHH. In particular, the ER-related modules and to a slightly lesser extent also modules related to other cellular injury response pathways are preserved in PHH. This suggests that, in general, the transcriptional programs that are driving cellular stress response pathways are maintained between primary liver hepatocytes and more de-differentiated models such as HepG2. Of interest, the cellular stress response gene networks are also well-preserved across species (human versus rat)^15^. The robust inter system-species preservation of the adaptive stress responses could be exploited to improve the translatability across test systems. The overall moderate preservation between HepG2 and PHH likely relates to the fact that HepG2 cells are transformed, de-differentiated and highly proliferative, requiring different wiring of transcriptional programs to drive HepG2 biology. However, since the PHH WGCNA was based on a larger number of substances, we cannot exclude that further refinement of the gene network organization could be achieved when more substances that impact of the biology of HepG2 cells would have been included. Regardless, we were able to define the conserved mechanisms between HepG2 and PHH, thereby delineating the biological applicability domain of the HepG2 test system to be used in a low tier test system approach that considers the HepG2 advantages in terms of cost, time, availability and robustness.

We have correlated HepG2 module EGs scores with cell death outcomes^17^ across the entire compound set, aiming to find common cellular responses leading to cellular adverse outcomes. This allowed us to identify the cellular responses that are activated to initiate repair and/or provide resilience to cell injury (adaptive mechanisms) and that are causally contributing to the adverse cell biological outcomes (adverse mechanisms). We revealed 11 modules with high positive correlation towards apoptotic and/or necrotic cell death of HepG2. A unique avenue in our current work has been to validate the association of these modules with cellular adverse outcomes by assessing the modulation of the cell death after nefazodone and nitrofurantoin treatment using RNA interference of selected genes. We sought a proof-of-concept and selected 67 genes that are contributing to cell signaling based on availability in a drugable siRNA genome library. We found five target genes that strongly reduced DILI compound-induced cell death upon the silencing *GTPBP2, TSC22D3, SIRT1, IRF1*, and *HSPA1B*. These results were indicative of the likely causal relationship between their upregulation and cell death. Of interest, these five genes are also modulated in PHH by various DILI compounds. Our RNA interference studies were based on individual gene depletion, thereby representing only a minor portion of the gene network. Thus, the causal contribution of the entire gene network is likely underestimated. One could foresee combined depletion experiments of multiple target genes or modulate the activity of transcription factors that are upstream of the individual gene network. Alternatively, CRISPR-Cas9 pooled library screens as they are applied to uncover resistance mechanisms against anticancer drugs, may open additional opportunities to more broadly map the causal-related genes of critical gene networks.

We further highlight the possible mechanisms of the five identified potential transcriptional key events that modulate DILI cell death. GTPBP2 is a family of GTP-binding proteins that have GTP hydrolase activity, and play an important role in cell signal transmission, cytoskeletal regulation, protein synthesis and other activities^45^. Although the mechanisms of GTPBP2 related to the hepatocellular injury is yet to be discovered, multiple studies have reported the involvement of GTPBP2 in ER stress^46,47^ and its interaction with ATF6^48^ reducing the activity of ATF6. The involvement of GTPBP2 in the ER stress is reflected by its parent module (HepG2:38) for ER stress-related responses. TSC22D3 or glucocorticoid-induced leucine zipper is associated with the glucocorticoid sensitivity also involved in the inflammation programs^48^. The upregulation of TSC22D3 was reported to be responsible for the cell death in cancerous cells^49^ possibly due to the activity of TSC22D3 to inhibit cell proliferation of the cancer cells. Thereby, the upregulation of *TSC22D3* might reduce the capacity of the cells to progress through the cell cycle and contribute to tissue regeneration. A previous study has attributed the effect of glucocorticoid on the activity of adenylate cyclase^50^. Specifically, the TSC22 protein family showed direct binding with this enzyme^51^. Moreover, the fact that *TSC22D3* is in WGNCA|HepG2:235 annotated for ‘adenylate cyclase-inhibiting activity’ suggested the connection between activity of this enzyme with proliferation^52^. SIRT1 (sirtuin1) is a member of class III histone deacetylase as a part of HepG2:31 annotated for ‘nuclear part’. The upregulation of SIRT1 in cells is found to be concurrent with NF-κB inflammation pathway-induced cell death^53^. IRF1 (interferon regulatory factor-1), a member of HepG2:41 annotated for ‘transcription factor’; IRF1 is a transcription factor regulating the gene expression during inflammation^54^. IRF1 has been reported to mediate liver damage during ischemia-reperfusion injury by activating the immune responses during the ischemia episodes^55,56^. HSPA1B is a member of heat shock protein 70 family and might play a role in the mechanisms of cell death related to DILI. *HSPA1B* is also a gene member of WGNCA|HepG2:64 which is annotated for ‘responses to heat’. Although heat shock protein families are known to repair the damaged cells^57^ (also showed by our results in Figure 5B – increase of cell death upon the perturbation of *HSPB1* and *HSPA8*), the upregulation of HSPA1B has been previously linked to the inflammation-induced cell death^58–60^. Although we managed to established these novel transcriptomics key events, further work needs to be performed to fully understand the mechanism of these genes in the course of DILI.

In conclusion, we have applied WGCNA to novel TempO-Seq toxicogenomics dataset. We anticipate that test system specific gene network-based approaches are powerful to efficiently mine the biology from toxicogenomics studies and contextualize the biology of the individual test systems through preservation statistics. Similar approaches would be worthwhile for other liver test systems that are relevant for DILI prediction, including HepaRG cells, iPSC-derived hepatocyte-like cells and liver microtissues. This would contribute to an improved understanding of the fit-for-purpose of individual liver test systems for mechanism-based safety assessment.

---

**Supplementary figure 1. Quality control output of the TempO-Seq RNA sequencing data**. (A) the library size of the samples grouped per compound from batch 1 (i) and batch 2 (ii). Each dot represent one samples. (B) The distribution of the CPM normalized count from batch 1 (i) and batch 2 (ii) grouped per compound. Each dot represent the read from one probe. (C) The correlation value (Pearson correlation) based on the CPM normalized count from batch 1 (i) and batch 2 (ii) of each samples to the mean value derived from the samples exposed to the same conditions (replicate). Each dot represent the correlation value from each sample. The color of the dots indicate the samples passing the threshold (Pearson correlation > 0.95 – blue : pass, red : not-pass).

**Supplementary figure 2. Transcriptomic analysis**. (A) The plot showing differential expressed genes number of FGF, HGF, and EGF at 8 hours (top) and 24 hours (bottom). Red bars indicate upregulated genes and blue bars indicate downregulated genes. (B) The correlation plot derived from the transcriptomic responses at the gene level (left), DEGs level (middle), and module responses (right) between nitrofurantoin (120 μM – 24 hours) from the batch 2 (y-axis) vs nitrofurantoin batch 1 (x-axis). Each dot shows the log2 fold change value of each gene (left) or module (right). The threshold of the DEGs is set with adjusted p-value < 0.01 and log2 fold change > [0.1]

**Supplementary figure 3. The weighted gene co-regulated network analysis applied to HepG2 transcriptomic data**. (A) The plot showing the pattern of scale free topology versus soft threshold power (left) and mean connectivity versus soft threshold (right). (B) The distribution of the size of the modules derived from the HepG2 transcriptomic data. (C) The overview of the gene memberships’ activities of the stress response modules upon the exposure of the positive compound at the particular time point and concentration (enlarged depiction). The box node indicate the hub gene (the gene with the highest correlation to the parent module). The color of the nodes represents the modulation and the size of the nodes represent the module correlation.

**Supplementary figure 4. WGCNA module activation by all tested substances**. (A) The modulation of the gene memberships of HepG2:38. The color of the plots represents the time point, the error bars indicate the standard error mean (SEM). (B) Heatmap showing the overview of the module responses. The heatmap contains 4 variables on the columns indicated by distinctive color groups: severity class of the compound, compound categories, time points, and dose level. These variables describe the exposure condition applied to the cells. Each row of the heatmap shows the eigengene scores of each module from every sample where red color shows activation and blue color shows repression. The column clustering of the heatmap is performed using the “*Ward d2*” algorithm applied to the Euclidian distance between aggregated variables (mean of the eigengene scores from all the dose levels and time points per compound). The row clustering of the heatmap is performed using the “*Ward d2*” algorithm applied to the Euclidian distance between modules.

**Supplementary figure 5. Module dynamics comparison between HepG2 and PHH**. (A) Distribution plots of log2 fold change values of gene memberships of HepG2:38, HepG2:64, and HepG2:65 grouped by time points in HepG2 and PHH upon the exposure of nitrofurantoin. (B) Module correlation plot between PHH and HepG2 exposed to nitrofurantoin (125 μM and 120 μM respectively) for 24 hours. Each dot on the correlation represents each module activity. The ER stress related modules are highlighted : HepG2:38, HepG2:64, HepG2:65. (C) (C) Response of HepG2 and PHH upon cyclosporine A exposure based on the HepG2:38, HepG2:64, and HepG2:65. The colors of the plot represent time points and the shapes of the dots indicate the cell lines.

**Supplementary figure 6. External trait-cell death correlation with the HepG2 gene networks**. (A) The density plot showing the density of the adjusted-p values of the correlation between the gene networks and cell death. The vertical line indicate the threshold for the significant correlation : adjusted p-value < 0.1. (B) The plots showing of the dynamics of 11 highest cell death-correlated modules upon the exposure of nitrofurantoin (i), troglitazone (ii), and diclofenac (iii). The colors of the plots represent time point.

**Supplementary figure 7.The expression measured in PHH of the genes whose perturbation reduced the cell death in HepG2 exposed to nitrofurantoin and nefazodone**. The heatmap shows the log2 fold changes of 18 out of 20 (2 genes are not measured in the TG-GATEs^12^) in PHH of the genes whose perturbation decreased the cell death fraction in HepG2 upon the exposure of nitrofurantoin (360 μM) and nefazodone (39.5 μM). The heatmap contains 3 variables on the columns indicated by distinctive color groups: severity class of the compound, time points, and dose level. These variables describe the exposure condition applied to the cells. Each row of the heatmap shows the log2 fold change values of each gene from every sample where red color shows upregulation and blue color shows downregulation. The column clustering of the heatmap is performed using the “*Ward d2*” algorithm applied to the Euclidian distance between aggregated variables (mean of the log2 fold change values from all the dose levels and time points per compound). The row clustering of the heatmap is performed using the “*Ward d2*” algorithm applied to the Euclidian distance between genes.

**Supplementary table – tab 1. List of the module memberships for each module Supplementary table – tab 2. Matrix containing the module scores of each experiment Supplementary table – tab 3. Annotations of the modules**

**Supplementary table – tab 4. SiRNA applied for gene silencing**

**Supplementary table – tab 5. Correlation values of the modules towards cell death**

**Supplementary table – tab 6. List of the high upregulated gene memberships of the high cell death-correlated modules**

**Supplementary table – tab 7. List of genes subjected to the RNAi experimentation with the description from the** http://www.genecards.org. These genes pass the threshold of adjusted p-val < 0.1 and log2 fold change > 2 at least in one condition.

**Supplementary table – tab 8. Z-scores value of the cell death upon perturbation of the target genes**

