## supplementary figure for "A Network-based Transcriptomic Landscape of HepG2 cells to Uncover Causal Gene Cytotoxicity Interactions Underlying Drug-Induced Liver Injury"

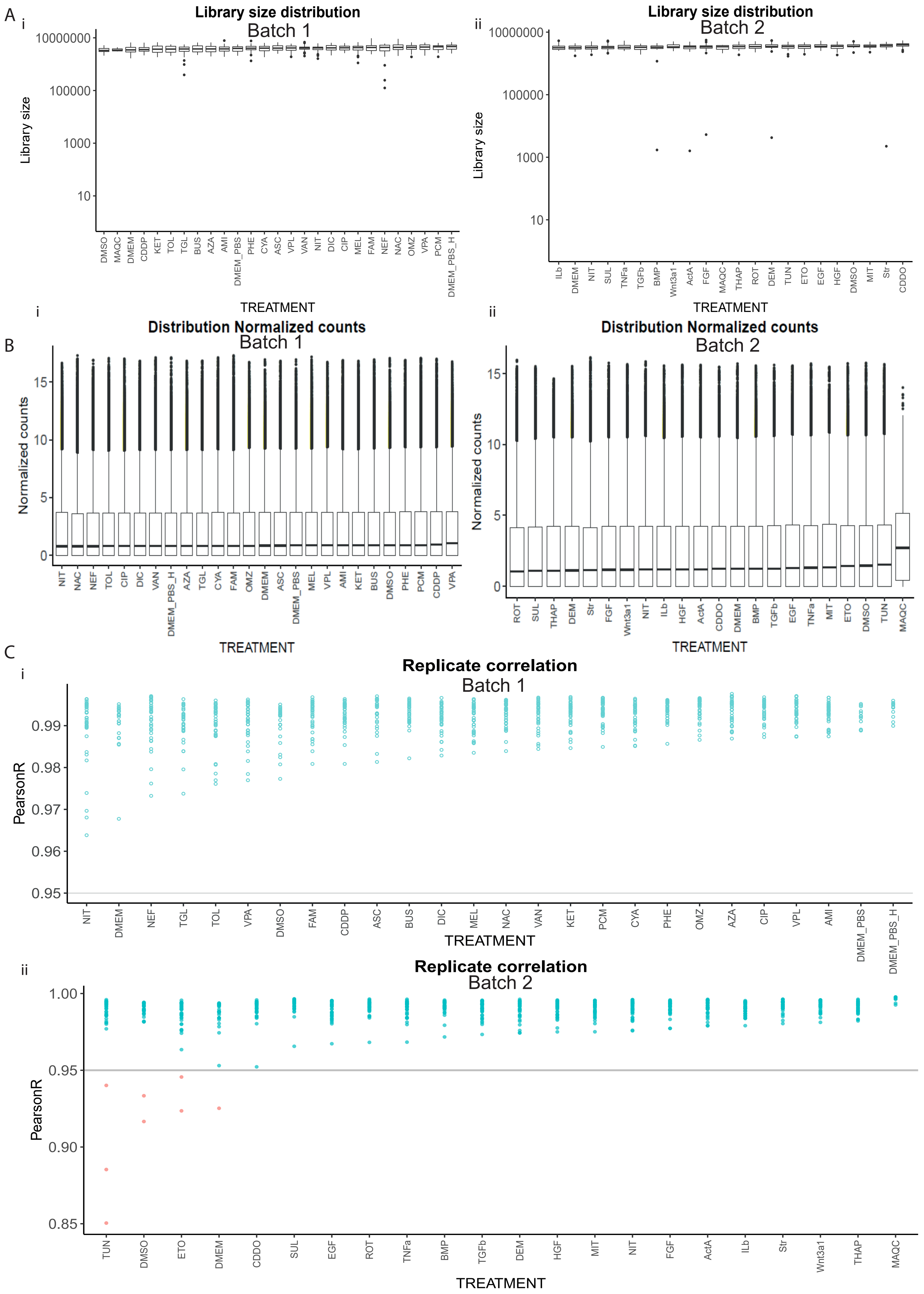


Supplementary figure 1. Quality control output of the TempO-Seq RNA sequencing data

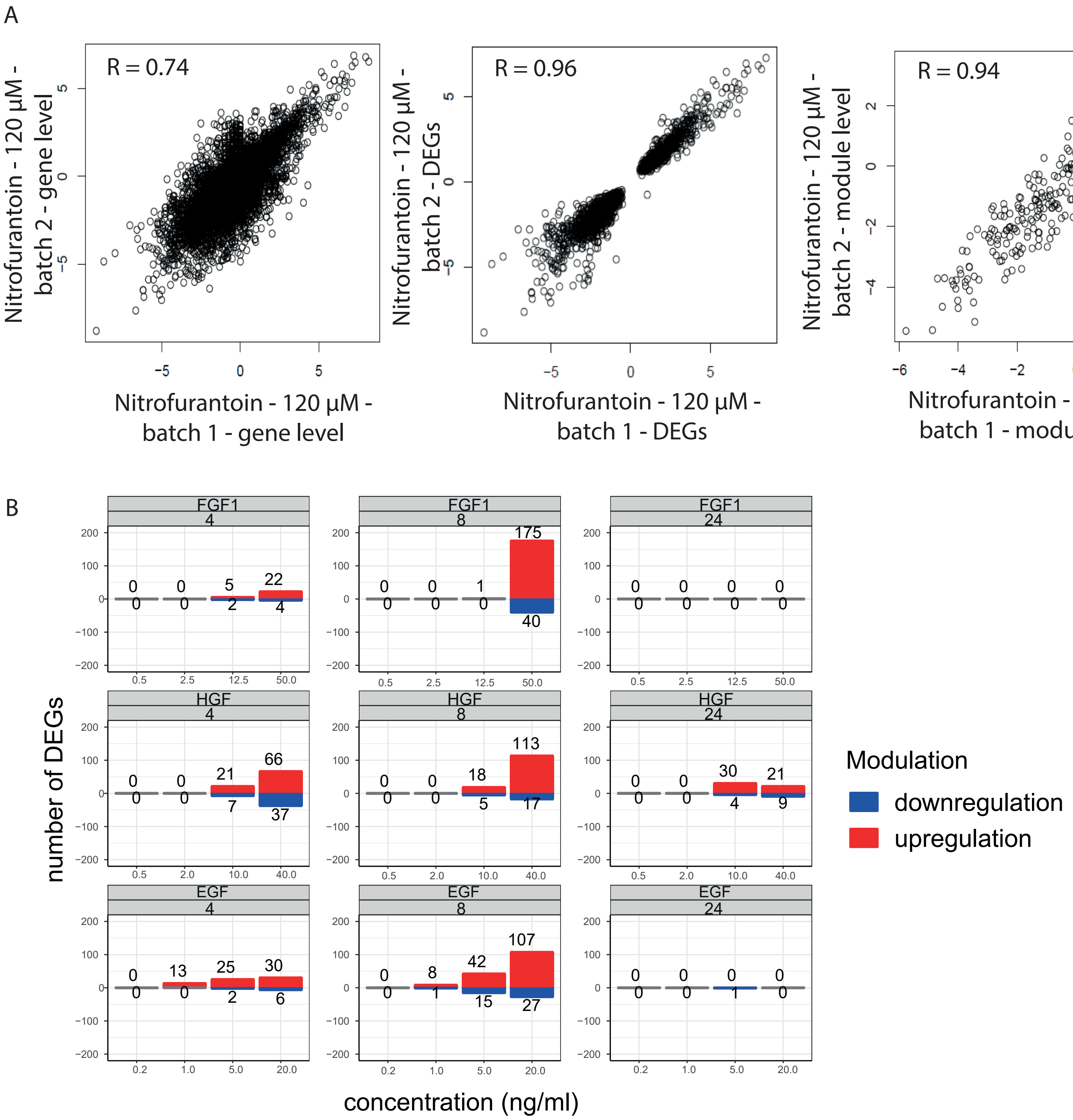


Supplementary figure 2. Transcriptomic analysis

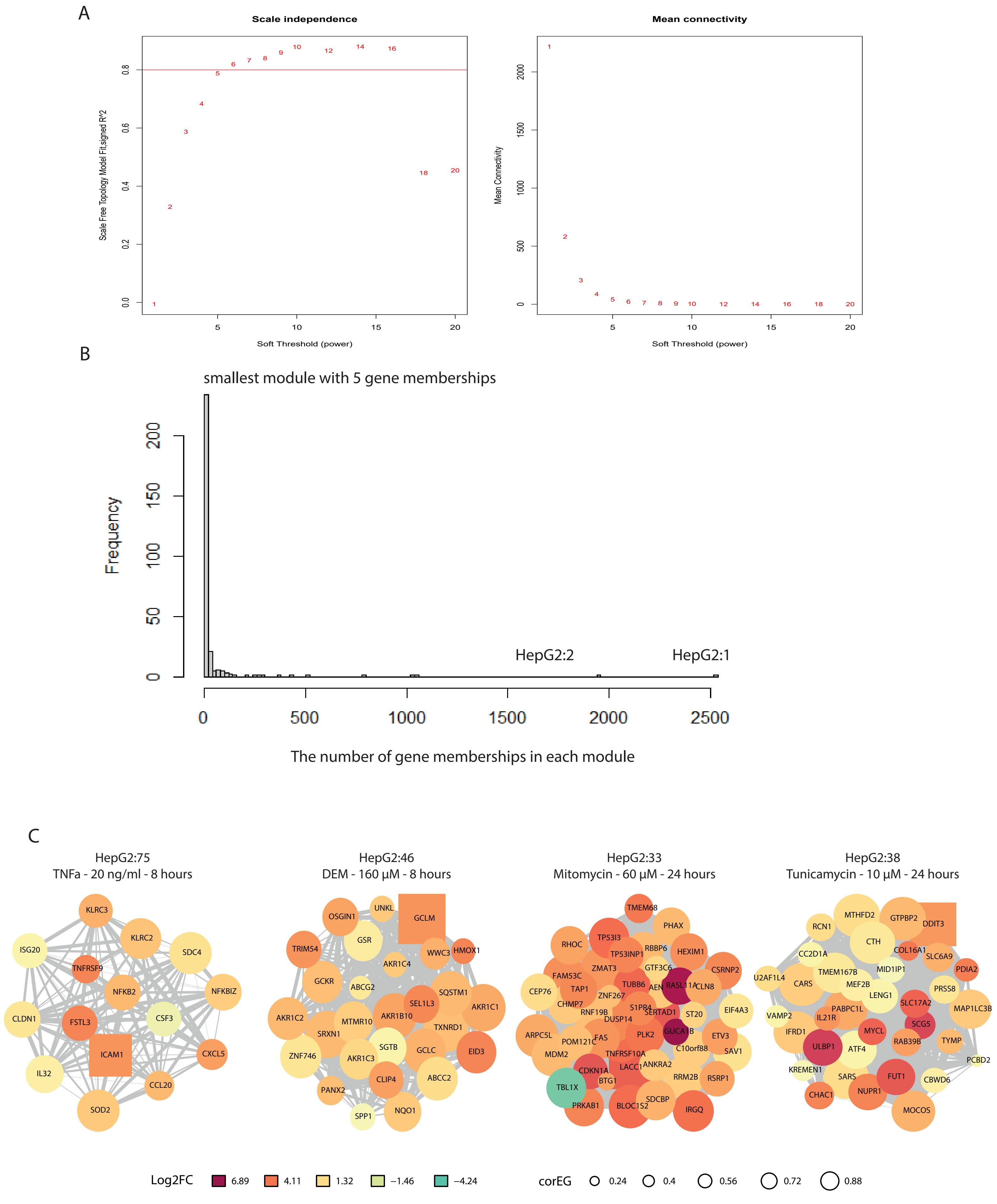


Supplementary figure 3. The weighted gene co-regulated network analysis applied to HepG2 transcriptomic data

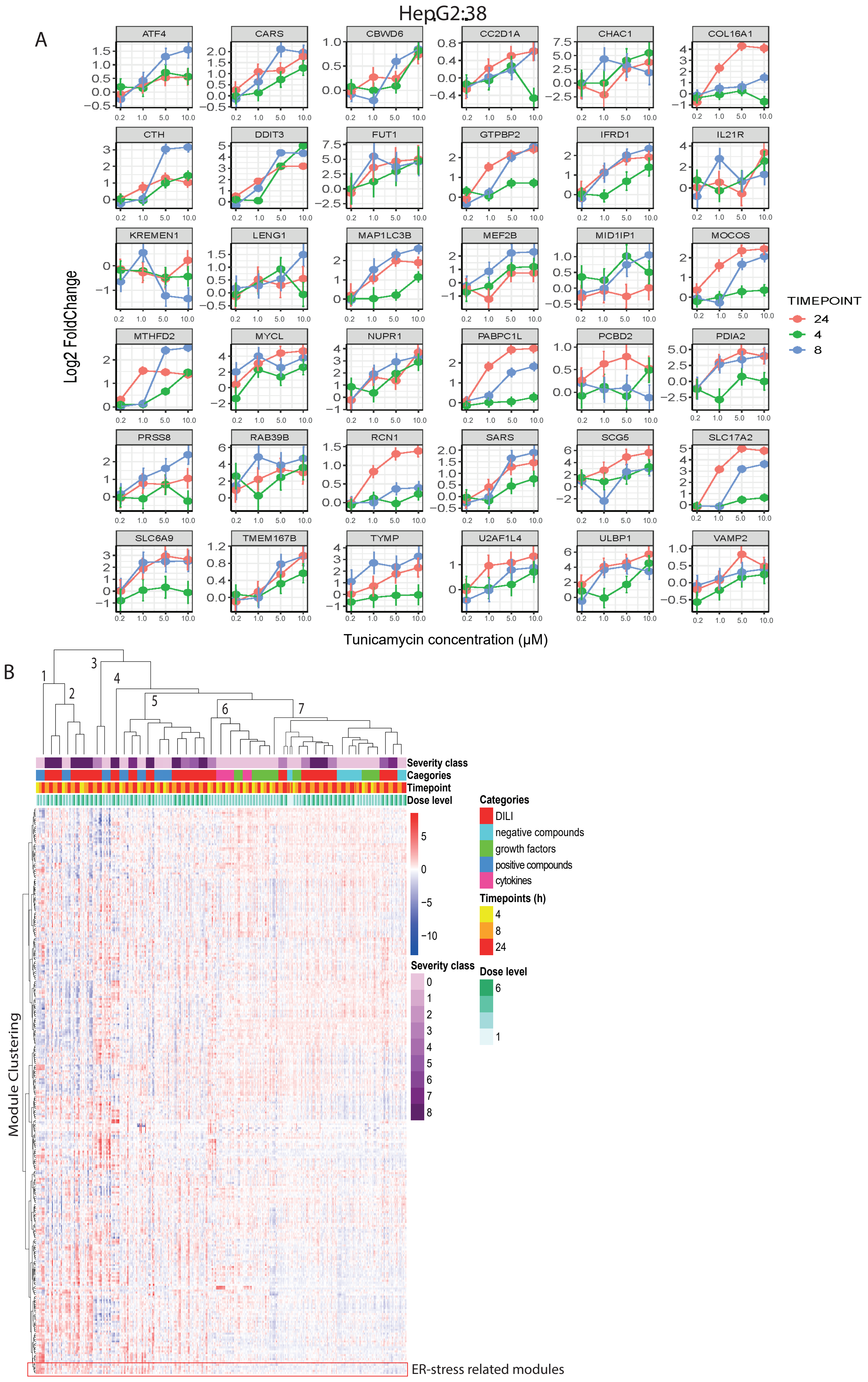


Supplementary figure 4. WGCNA module activation by all tested substances

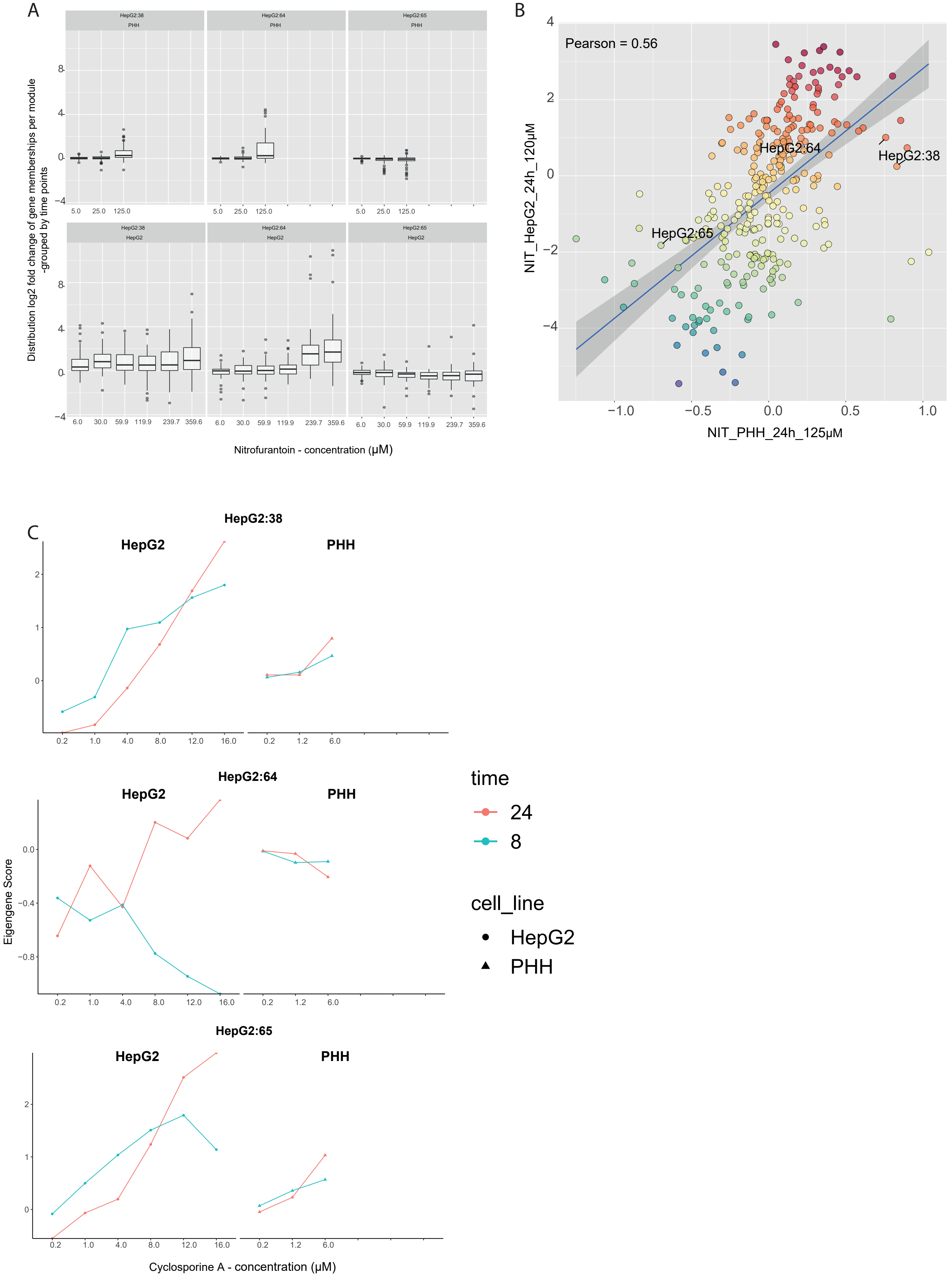


Supplementary figure 5. Module dynamics comparison between HepG2 and PHH

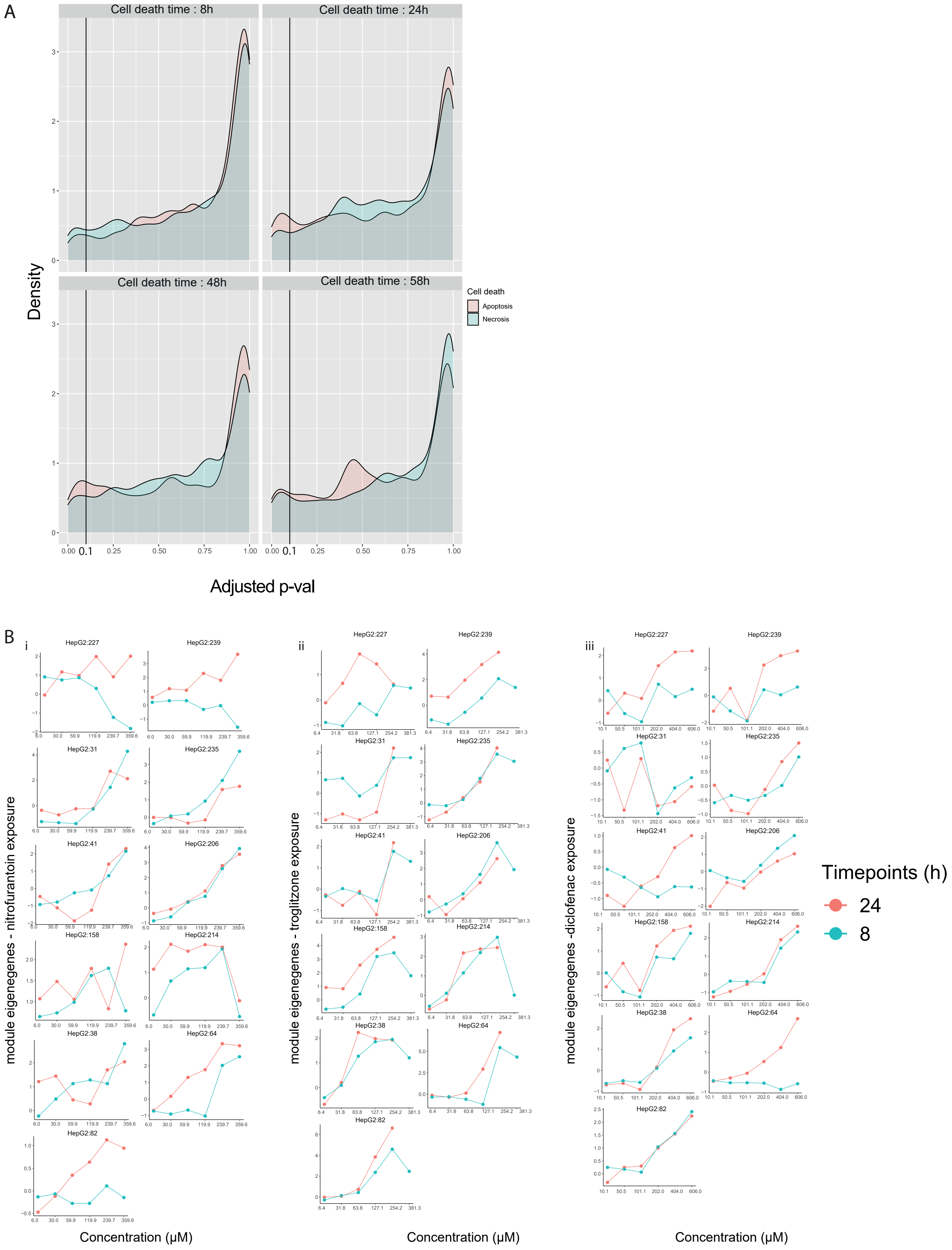


Supplementary figure 6. External trait-cell death correlation with the HepG2 gene networks

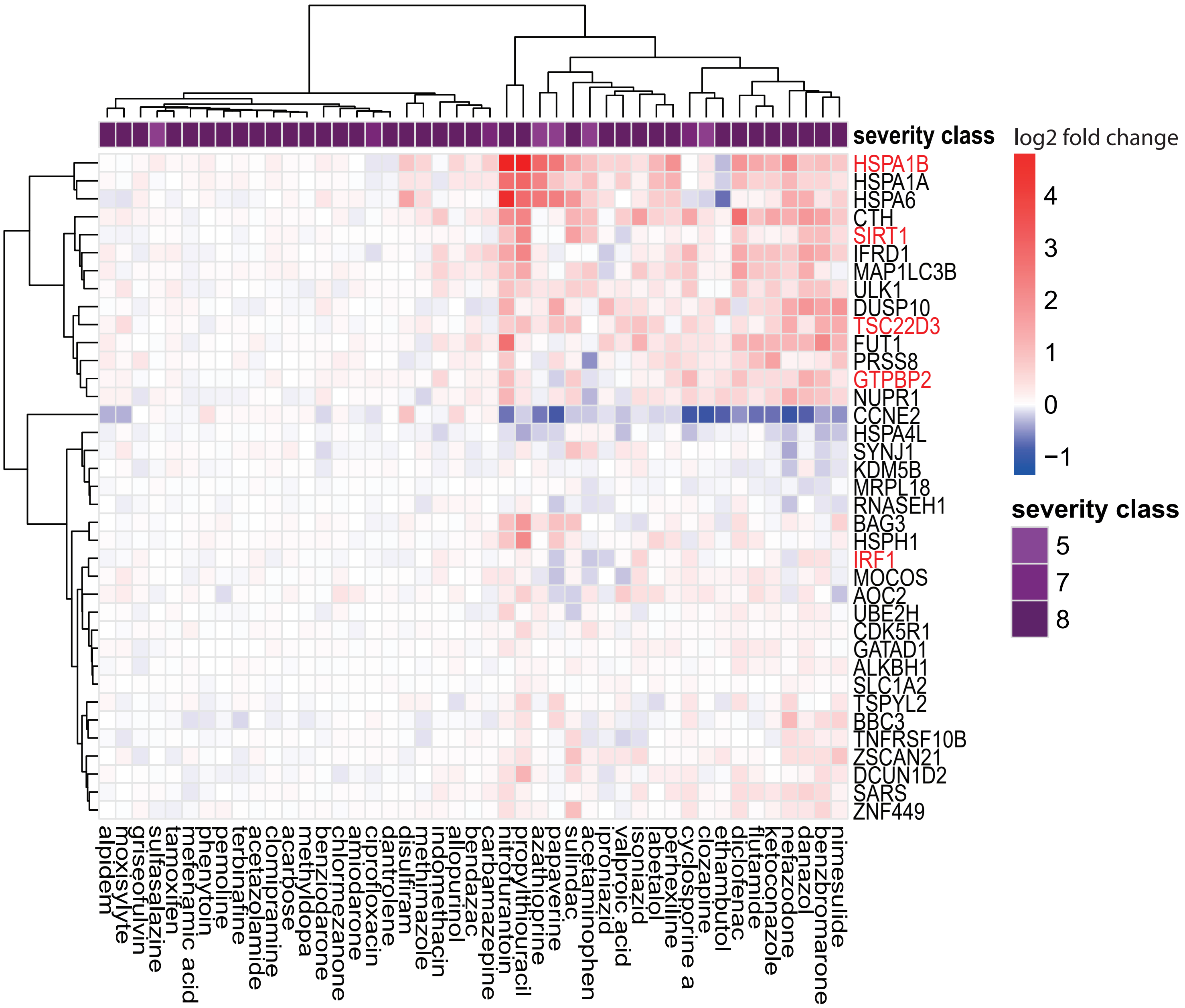


Supplementary figure 7.The expression measured in PHH of the genes whose perturbation reduced the cell death in HepG2 exposed to nitrofurantoin and nefazodone
